# The active secretion of a subunit of IL-12 by tissue cells is regulated by Valosin-Containing Protein and intracellular calcium redistribution

**DOI:** 10.64898/2026.01.28.702376

**Authors:** Elizabeth M. Hill, Jaclyn Bredenkamp, Gabrielle C. McBroom, Budhaditya Chatterjee, Kenneth M. Rosenberg, Nevil J. Singh

## Abstract

One molecular mechanism for collaboration between innate immune and tissue stromal cells in regulating effector responses is the formation of the innate cytokine IL-12 as a result of the two cell types contributing distinct subunits (p40 and p35) allowing for the localized assembly of the functional IL-12 heterodimer. Dendritic cell activation leads to p40 secretion, but the pathways in stromal cells allowing p35 secretion are unknown. We identify the inhibition of Valosin-Containing Protein (VCP) and increases in intracellular calcium signaling as the key regulators for the secretion of p35 from its homeostatic reservoirs in the endoplasmic reticulum. This p35 secretion, in the absence of co-expression of the p40 subunit, requires active gene expression, is independent of ER stress pathways, and is distinct from passive (cell-death dependent) release processes. Thus, an active sensory apparatus in non-hematopoietic cells contributes to the collaborative control of the trajectory of early immune differentiation.

**One Sentence Summary:** VCP inhibition and calcium flux allows stromal cells to release the p35 subunit of the cytokine IL-12 and regulate localized cytokine activity.

## INTRODUCTION

A typical T cell response involves the differentiation of naïve cells to acquire effector functions that are specifically tailored to the type of threat faced by the organism as well as the tissue that is affected. Canonically, antigen-presenting dendritic cells (DC) are known to play an instructive role in this process by secreting innate cytokines. For instance, DCs secreting IL-12 vs IL-23 enhance the differentiation of IFN-γ vs IL-17 producing effector T cells respectively, which have distinct roles in combating intracellular vs extracellular pathogens. It is increasingly clear, however, that non-hematopoietic, tissue cells also participate in shaping the trajectory of the immune response (*1–3*), although the underlying mechanisms are less clear. In this context, we have previously shown (*4*) that stromal cells and immune cells can molecularly collaborate to produce functional IL-12 *in vivo* leading to maintenance of tissue-specific IFN-γ-producing T cells during a *Leishmania* infection.

The logic for this partnership derives from the unique biology of the IL-12 family of cytokines. IL-12 is a heterodimeric cytokine that typically forms by the covalent association of α and β subunits also known as p35 (IL-12p35, IL-12α) and p40 (IL-12p40, IL-12β) (*5, 6*). Innate immune cells (mostly macrophages and DCs) secrete IL-12 when they are appropriately stimulated – usually by microbial products or alarmins through activation of pattern recognition receptors (PRR) signals in the early phases of the immune response, in combination with accessory signals such as CD40L or IFN-γ itself (*7*). Molecularly, the p35 and p40 subunits of IL-12 are made within the myeloid cells and linked covalently by a disulfide bond before being secreted as a heterodimer. However, in addition to the secretion of this fully assembled cytokine by activated myeloid cells, we and others have shown that copious amounts of just the p40 monomer is secreted early in the response (*8, 9*). In contrast, the p35 subunit of IL-12 is not freely secreted as a monomer and typically requires the p40 subunit to be co-transcribed in the same cell for its proper folding and secretion (*10*).

Despite this, the p35 transcript is reportedly expressed in many non-immune cell types that do not co-express p40, although an IL-12 independent function for p35 in these cells is yet to be described (Human Protein Atlas; https://www.proteinatlas.org/ENSG00000168811-IL12A/single+cell+type). Some of this is accounted for by the less characterized cytokine IL-35, which is formed by the association of p35 and EBI3; but EBI3 expression is also mostly limited to lymphoid and myeloid lineages (*11–13*). Therefore, it is reported that in the absence of p40, p35 forms misfolded homodimers that are subject to ER-associated degradation (ERAD) (*10*). Within the ER, p35 also interacts with a variety of resident chaperone proteins including BiP, calreticulin, and protein disulfide isomerases, many of which are considered critical for its retention in the absence of p40 (*14*). Despite this stringent compartmental regulation of p35 formation and IL-12 assembly, we have previously shown that p35 released from cells after lytic freeze/thaw can combine with p40 collected from serum, to generate functional IL-12 in vitro (*15*). Validating this paradigm in an *in vivo* setting, when the subunits of IL-12 are separated into two different cellular sources using a bone marrow chimera model, the IFN-γ response to *Leishmania* was still maintained (*4*). These results suggested that the two subunits of IL-12 can be released and then combined extracellularly to generate the biologically active cytokine. Importantly, this mode of IL-12 production (which we previously termed as two-cell IL-12) relies on the ability of p35 to be released from cells where p40 is not expressed, despite the restrictions (discussed above) that are canonically placed on p35 being secreted independently. As a cellular mechanism for coordinating responses between stromal cells and innate immune cells this also offers the potential for two independent sensing mechanisms to trigger IL-12 formation towards affecting T cell differentiation (*1*).

The mechanisms of p40-independent secretion of p35 and importantly, the triggers in stromal cells that stimulate such pathways are not known. Our previous studies suggested that in the context of an ongoing response, the release of p35 may be passive – i.e. a consequence of lytic cell death of the cell expressing p35, following infection or inflammation. This hypothesis is consistent with other well-established findings showing that the induction of cell death or damage can lead to the release of proteins (including DAMPs, alarmins) that are usually nuclear- or ER-restricted (*16–18*), often leading to immune activation. Intriguingly, using sensitive reporter systems for tracking p35 release from non-hematopoietic cell lineages, we find that pharmacological activation of cell death led to only a minimal release of p35, even under conditions where a control protein was passively released from dying cells. This led us to investigate the possibility that p35 is released from non-hematopoietic cells actively – in response to microbial or cell-health indicators, to promote the two-cell formation of IL-12. To identify the underlying mechanisms of this process, we screened a library of compounds affecting ER-related functions in cells. Successful “p35-releasers” did not cluster with a specific landscape of cellular functions but implicate intracellular cationic balance as well as the pleiotropic regulator Valosin-Containing Protein (VCP) as new targets. During disruption of either pathway, p35 appears to be freed from ER residency and translocate through the Golgi, albeit with different kinetics. p35 release not completely reliant on the activation of canonical stress signaling pathways such as the unfolded protein response (UPR). Finally, CB-5083, but not ionomycin, required active gene expression to trigger p35 release – leading us to a new model for regulated stromal release of p35 without p40 co-expression (Figure 8J). Together, these results implicate a previously unknown sensory apparatus at the nexus of intracellular calcium and VCP function that regulates the active release of p35 from viable non-hematopoietic cells. This paradigm can lead to a revision of current models of immune differentiation and the expansion of targets for ‘adjuvanting’ the immune response in local tissue microenvironments.

## RESULTS

### The IL-12p35 subunit is not secreted in the absence of the IL-12p40 binding partner

The p35 subunit of IL-12 is rarely secreted by itself in healthy cells and measuring the extracellular amounts of this protein has been challenging, largely due to the paucity of robust reagents (*10*). Therefore, we developed sensitive luciferase- and fluorescence-based reporters to measure the trafficking and potential release of IL-12p35 in cells by fusing p35 cDNA with the NanoLuc luciferase (p35NL) or the mScarlet fluorescent protein (p35-Scarlet). In the canonical pathway, p35 is not released from cells unless p40 is also co-expressed. Accordingly, we validated this relationship by measuring luminescence in the culture medium from the p35NL reporter in the presence or absence of p40. p35NL is retained in cells when expressed on its own, like native p35 (Figure 1A, 2.0µg/mL light red bars, are not significantly different from NanoLuc-only controls, light blue bars, p = 0.1164, unpaired t-test). When increasing concentrations of the p35NL construct were co-transfected with p40, increased luminescence in the cell culture supernatant indicated p35NL release (Figure 1A, red bars). For instance, at the 2.0µg/mL dose of p35NL plasmid (Figure 1A) there was a significant increase in luminescence in the supernatant of p35NL + p40 compared to p35NL + EV (p < 0.0001, unpaired t-test). In experiments with a control NanoLuc-only (NLO) construct, no significant increase (p = 0.4757, unpaired t-test) in luminescence in the supernatant was observed with p40 co-transfection (Figure 1A, dark blue bars). Together, these results validate that p35NL is dependent on p40 for its release like native p35 and that the p35NL luciferase assay provides a sensitive method to quantify the release of the p35 subunit into the supernatant.

**Figure 1.**
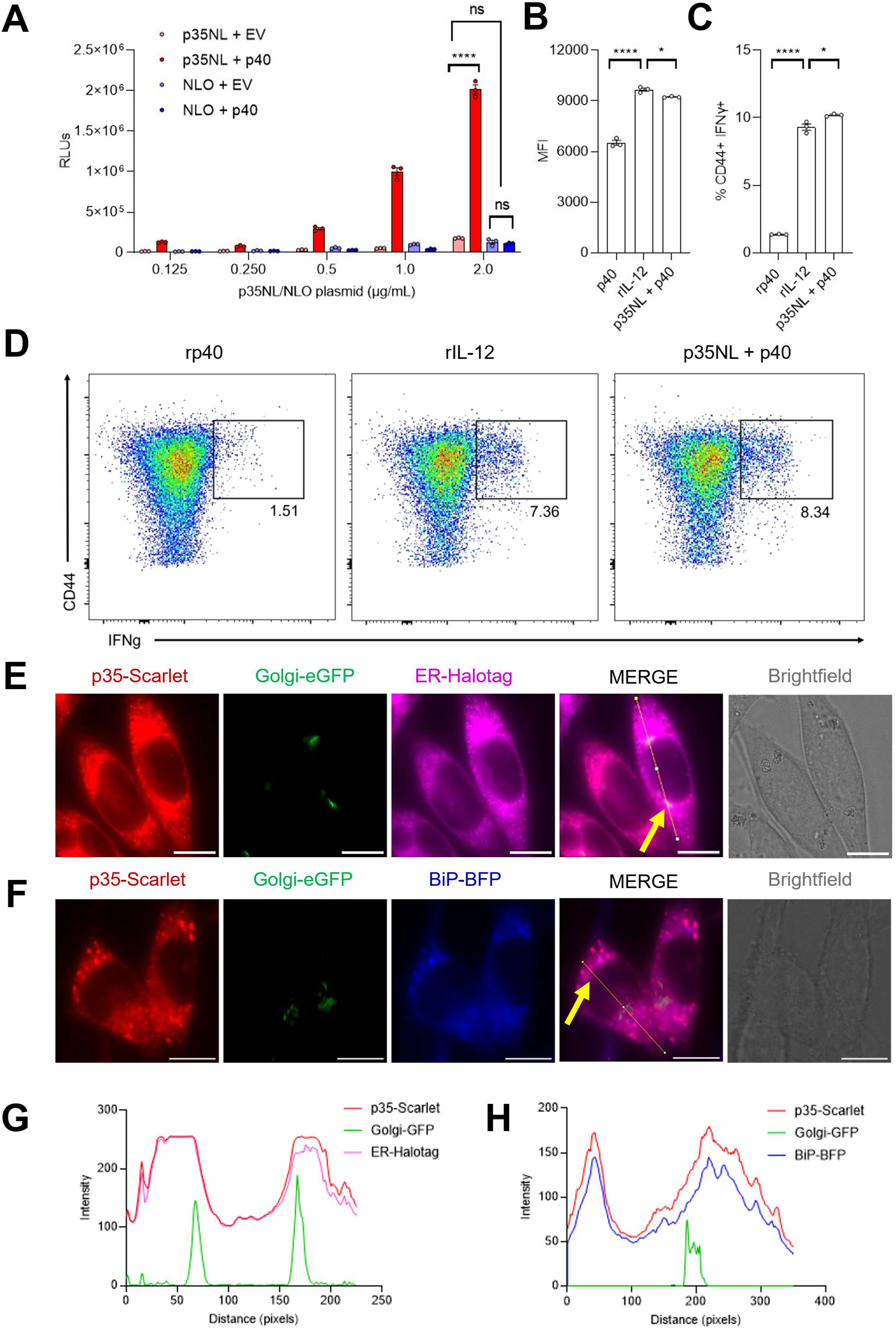
The IL-12p35 subunit is not secreted in the absence of the IL-12p40 binding partner: Development of two novel reporter systems to track the release of p35 from cells and subcellular location throughout p40-independent release. **(A)** Luciferase assay to detect p35NL release in the cell culture supernatant when co-expressed with p40. HEK cells were transfected with p35NL + empty vector (light red), p35NL + p40 (red), NanoLuc-only (NLO) + empty vector (light blue), or NLO + p40 (blue). Supernatants were collected 48 hours after transfection for luciferase assay (n=3). Luminescence data is represented in Raw Luminescence Units (RLUs). Functionality of p35NL as an IL-12 subunit when co-expressed with p40. SMARTA transgenic T cells were activated in vitro and re-stimulated with either recombinant p40 protein, recombinant IL-12, or supernatants from HEK cells co-transfected with p35NL and p40. After 2 hours of re-stimulation, IFN-γ production by T cells was measured using intracellular cytokine staining. **(B)** MFI of live CD4^+^CD44^+^IFN-γ^+^ SMARTA transgenic T cells populations after restimulation (n = 3). **(C)** Percentages of MFI of live SMARTA CD4^+^CD44^+^IFN-γ^+^ transgenic T cells populations after restimulation (n = 3). **(D)** Representative flow plots compare the percentage of live CD4+CD44+IFNγ+ cells among groups (n = 3). Characterization of fluorescent reporter system for tracking the intracellular location of p35 through ER and Golgi compartments. **(E)** Murine fibroblasts (L cells) were retrovirally transduced with p35-Scarlet, Golgi-eGFP (Beta-1,4-galactosyl transferase 1), and ER-Halotag (KDEL tagged with Halotag and developed with JFX650). Images are representative of steady-state cells during live cell fluorescent imaging at 100X. White scale bars correspond to 10µm. Characterization of fluorescent reporter system for tracking the association of p35 with ER chaperone, BiP. **(F)** L cells were retrovirally transduced with p35-Scarlet, Golgi-eGFP, and BiP-BFP. Images are representative of steady-state cells during live cell fluorescent imaging at 100X. White scale bars correspond to 10µm. **(G)** Pixel intensity of each marker in p35-Scarlet/Golgi-GFP/ER-Halotag L cells along the ROI line draw in ‘MERGED (F)’. ROI line is designated by an arrow. (H) Pixel intensity of each marker in p35-Scar/Golgi-GFP/BiP-BFP L cells along the ROI line draw in ‘MERGED (G)’. ROI line is designated by an arrow. Data is presented as the mean with SEM error bars. Significant changes in luminescence in (A) were determined using an unpaired t-test. Significant changes in percentage and MFI in (B) and (C) were determined using an unpaired t-test. p < 0.05; ** p < 0.01; *** p < 0.001; **** p < 0.0001; ns = not significant.

We also confirmed that p35NL protein released with the p40 subunit is biologically active, by evaluating the ability of supernatants produced in Figure 1A to enhance IFN-γ production in activated T cells as IL-12 is known to do (Figure 1B, C, D). When *in vitro* activated SMARTA transgenic T cells specific to the LCMV GP_61-80_ peptide were re-stimulated in the presence of the supernatants from p35NL + p40 transfected cells, the production of IFN-γ by activated SMARTA T cells was significantly increased compared to treatment with recombinant p40 alone (p < 0.0001, unpaired t-test). The percentages of CD44+IFN-γ+ T cells triggered by p35NL + p40 were similar to those in recombinant IL-12 treated cultures (Figure 1C, p = 0.0229, unpaired t-test). These results confirm that p35NL is a functional IL-12 subunit, capable of pairing with p40 to signal and stimulate IFN-γ by T cells.

To track the trafficking of p35 throughout subcellular compartments during release in the absence of p40, we also generated a fluorescent reporter in which the murine p35 protein is linked to the red fluorescent protein, mScarlet, (as well as a Myc tag). Using live-cell, widefield fluorescent microscopy, L cells retrovirally transduced with p35-Scarlet showed a cytoplasmic localization of the protein, consistent with other reports finding p35 in foci throughout the cytoplasm when expressed in the absence of p40 (*19*). We retrovirally transduced L cells expressing p35-Scarlet with additional fluorescent markers for different subcellular compartments: Golgi-eGFP (which marks the trans-Golgi through beta1,4-galactosyltransferase 1) and ER-Halotag (which marks the endoplasmic reticulum by virtue of a KDEL sequence). Tracking p35-Scarlet, Golgi-eGFP and ER-Halotag allowed us to determine the sub-cellular localization of p35-Scarlet in the steady state. p35-Scarlet overlapped with the localization of ER-Halotag (Figure 1E, G) with very little overlap with Golgi-eGFP. Finally, we used a fluorescently tagged version of the ER chaperone, BiP (BiP-BFP) to show that p35 co-localizes with BiP in steady-state stromal cells (Figure 1F, H). These results validate an extensive palette of reporters that recapitulate the biology of p35 protein in non-hematopoietic lineage cells, helps track its localization in the ER through live cell fluorescent imaging, and can sensitively measure its secretion – which is not evident in the absence of p40 expression.

### Cell death is not a major driver of p40-independent p35 release from non-immune cells

As discussed in the introduction, the impetus for this study is our previous finding that p35 can be released from non-hematopoietic cells in a mouse model of *Leishmania* infection and then drive protective IFN-γ responses by associating with p40 in the extracellular milieu to form bioactive IL-12 (*4*). We initially hypothesized that pathogen-induced cell death could lead to p35 release from ER-trapped pools in a manner previously reported for alarmins (immunogenic DAMPs such as ATP and HMGB-1) or pro-inflammatory cytokines (e.g., IL-1β and IL-33). To test this hypothesis, the murine fibroblast L292 cell line (L cells) was stably transduced with p35NL or NLO to generate reporter cells that could be treated with various pharmacological inducers of cell death (Figure 2). First, we tested actinomycin D, a DNA-intercalator that can cause cell death at high doses (*20*). Accordingly, after treatment with actinomycin D, viability of both p35NL and NLO cells was significantly reduced at doses greater than 5µg/mL (Figure 2A right, compared to DMSO, p = 0.0047) (NLO – Figure 2B right, p = 0.019, one-way ANOVA). At this dose, there was an ∼10-fold increase in supernatant RLUs in the NLO cells compared to DMSO controls, indicating that intracellular NanoLuc was released passively upon cell death (Figure 2B). There was increased p35NL in the supernatant as well (p35NL – Figure 2A left, compared to DMSO, p < 0.0001) but this was much more modest ∼2-fold increase (Figure 2C) suggesting that passive release of p35 was more restrained, compared to NLO control protein.

**Figure 2.**
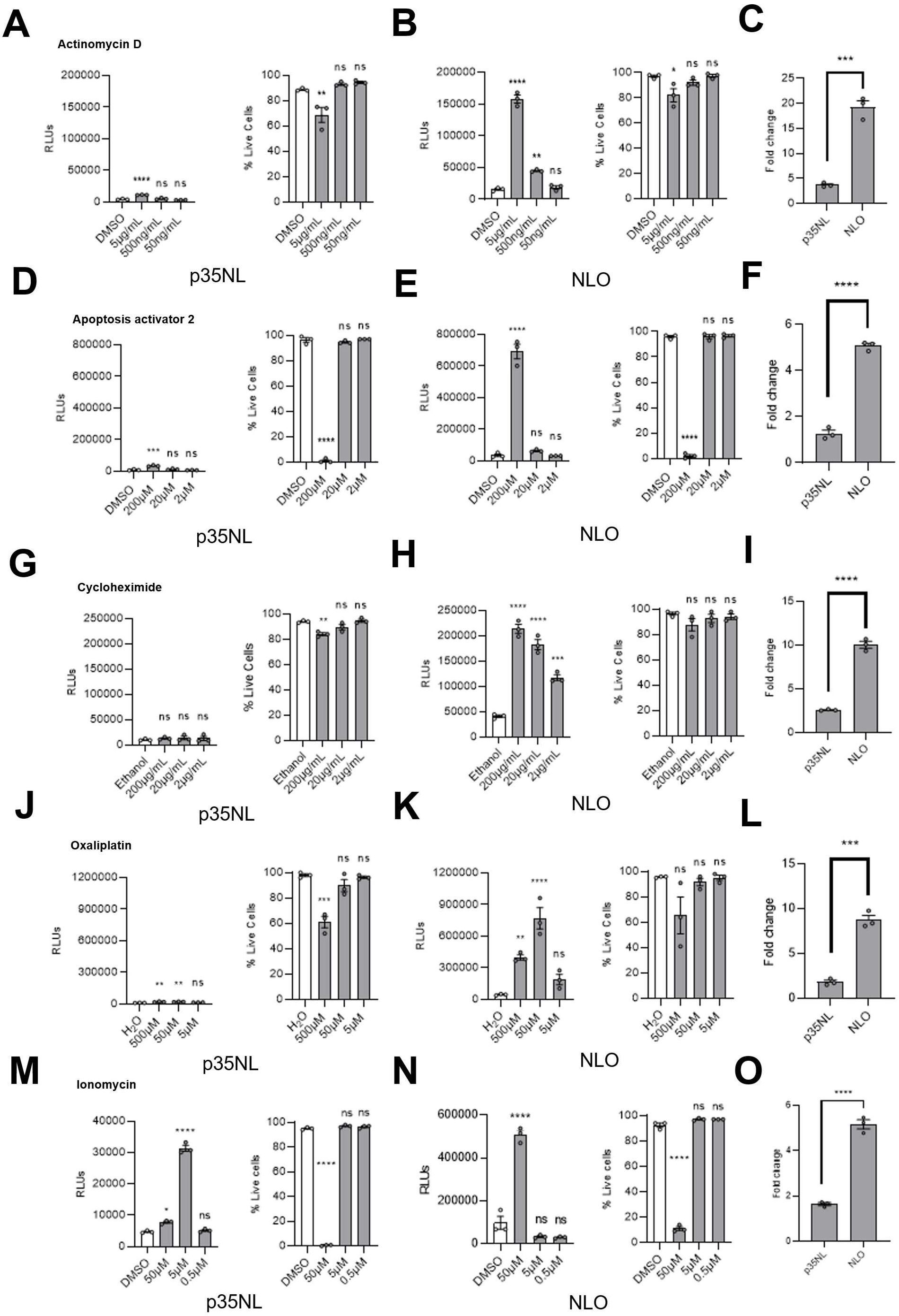
Cell death is not a major driver of p40-independent p35 release from non-immune cells. Murine L cells were retrovirally transduced with p35NL or NLO constructs. Cells were exposed to several chemical inducers of cell death. Supernatants were collected for luciferase assay after 24 hours of exposure to drug (n=3). Cell viability after 24 hours of drug exposure was measured using propidium iodide staining (n=3). Luciferase assay of supernatants from p35NL cells exposed to (**A**, left) actinomycin D, (**D**, left) apoptosis activator 2, (**G**, left) cycloheximide, (**J**, left) oxaliplatin, and ionomycin (**M**, left). Viability of p35NL cells after exposure to (**A**, right) actinomycin D, (**D**, right) apoptosis activator 2, (**G**, right) cycloheximide, (**J**, right) oxaliplatin, and ionomycin (**M**, right). Luciferase assay of supernatants from NLO L cells exposed to (**B**, left) actinomycin D, (**E**, left) apoptosis activator 2, (**H**, left) cycloheximide, (**K**, left) oxaliplatin, and ionomycin (**N**, left). Viability of NLO cells after exposure to (**B**, right) actinomycin D, (**E**, right) apoptosis activator 2, (**H**, right) cycloheximide, (**K**, right) oxaliplatin, and ionomycin (**N**, right). Fold change of luminescence comparing vehicle controls (DMSO, Ethanol, and H_2_O) to the highest dose of drugs from cells treated with **(C)** actinomycin D, **(F)** apoptosis activator 2, **(I)** cycloheximide, **(L)** oxaliplatin, and **(O)** ionomycin. Data is represented as the mean raw Relative Luminescence Units with SEM error bars. Significant elevations in luminescence were determined through ordinary one-way ANOVA of the mean in comparison to vehicle controls (DMSO, Ethanol, and H_2_O) Significant changes in cell viability were determined through an ordinary one-way ANOVA of the mean in comparison to vehicle controls (DMSO, Ethanol, and H_2_O). *, p < 0.05; ** p < 0.01; *** p < 0.001; **** p < 0.0001; ns = not significant

In response to treatment with apoptosis activator 2 (a broad stimulator of caspases), both cell lines were not viable at doses of 200µM (p35NL – Figure 2D right, compared to DMSO p < 0.0001) (NLO – Figure 2E right, compared to DMSO, p < 0.0001). Again, while cell death did lead to detection of p35NL and NLO in the cell culture supernatant at 200µM AA2 (p35NL – Figure 2D left, compared to DMSO, p < 0.0001) (NLO – Figure 2E left, compared to DMSO, p < 0.0001), the release was considerably higher in NLO cells (Figure 2F). A third representative cytotoxic agent, high doses of the translation inhibitor cycloheximide, viability was significantly reduced in p35NL cells (Figure 2G right, compared to DMSO, p = 0.0023). Despite the decrease in viability, cycloheximide did not significantly induce release of p35NL (Figure 2G left, compared to DMSO, p > 0.99); however, there was a 5-fold, increase for NLO cells (Figure 2H, I, compared to DMSO, p < 0.0001).

Finally, we evaluated oxaliplatin, a chemotherapy agent known to induce cell death through DNA damage. Additionally, oxaliplatin has been linked to the induction of “immunogenic cell death” and the release/exposure of DAMPs such as calreticulin (*17*). Viability of p35NL cells was significantly reduced (dose of 500µM in Figure 2J, compared to DMSO, p = 0.0002). Viability was reduced in NLO cells in response to the highest dose of oxaliplatin but not significantly (Figure 2K right, compared to DMSO, p = 0.0616). As with the previous cytotoxic agents, oxaliplatin also effectively released NLO but less robustly p35NL (p35NL – Figure 2J left, compared to DMSO, p = 0.0044) (NLO – Figure 2K left, compared to DMSO, p = 0.0072, F3,8 = 30.16). Interestingly, at doses of oxaliplatin where L cells were still viable (50µM) both p35NL and NLO were still released (p35NL – Figure 2J left, compared to DMSO, p < 0.0001) (NLO – Figure 2K left, compared to DMSO, p = 0.0072) suggesting that p35NL and NLO release could precede when apoptotic cells become positive for propidium iodide.

In stark contrast to these data, the treatment of cells with the calcium ionophore, ionomycin, which has also been reported to trigger the release of immunogenic DAMPs such as HMGB1 at high doses (*18*), produced an intriguing pattern. While it did not cause p35NL release at low or high doses (Figure 2M, left), the intermediate (5μM) dose led to a significant increase in p35NL release that was not present in the NLO cell line (Figure 2N, left). At the 5µM dose, however, the cells were viable (Figure 2M, right) but significant cell death occurred at 50µM. In contrast to the p35NL release, NLO was retained in cells until viability was lost (p35NL – Figure 2M, p < 0.0001) (Figure 2O, left).

As an additional measure of cytotoxicity and lysis of cells with chemical inducers of cell death, we quantified LDH release (Supplemental Figure 2). While the overarching hypothesis prior to this work (*4*) was that cell death was a critical trigger of p35 release from non-immune cells, the data in this section suggest that such a lytic release was not the most efficient mechanism for p35NL release compared to NLO (see also Figure 4 for additional visualizations of this data). Importantly, we find that it was also possible to have p40-independent release of p35 from viable cells by treatment with at least one drug, the Ca^2+^ ionophore ionomycin.

### A pharmacological screen reveals key cellular pathways regulating p40-independent p35 release

Based on the activity demonstrated by ionomycin, we developed a high-throughput pharmacological screen to map cellular pathways involved in p40-independent release of p35 during inflammation and in the context of two-cell IL-12. We curated a library of 115 (including ionomycin) drugs that are specifically known to target ER retention, ionic balance, innate immune stimulators, ER stress, unfolded protein response activators, and other specific modes of cell death (Figure 3, rows represent individual drugs and columns indicate biological pathways they target). p35NL cells were exposed to titrations of each drug ranging from 100-0.09μM for 24 hours, and the supernatant was probed for p35NL luminescence. We used a threshold criterion for inducers of p35 release where a fold change of >4 compared to untreated p35NL cells was considered a “positive inducer”. Using these criteria, we identified 8 pharmacological inducers of p35 release (Figure 3A, Supplemental Figure 2): BiP inducer X, BRITE33873, Calmidazolium chloride, BHQ, KB-R7943 mesylate, Golgicide A, CB-5083, and Ionomycin. These 8 drugs displayed different dynamics regarding how they induce p35 release. Some drugs only released p35 at the highest doses (Supplemental Figure 2C, 2F); most likely at the doses where they are lethal. 93% of drugs that were screened had no effect on p35NL release, emphasizing that rather than a broad dysregulation of ER functions, at least one distinct pathway (reliant on VCP) is critical for p40-independent p35 release. While pharmacological agents can vary quite a bit in their target-specificity, we used available information about these compounds to perform unbiased hierarchical clustering of the activities of our library (Figure 3B). Based on the literature, while some of these drugs formed clusters with related functional activities (e.g. top left cluster, showing a convergence in their ability to trigger Apoptosis, ROS increase, as well as activating the UPR/ER Stress pathways), none of these clusters corresponded to the release of p35 (Figure 3B) This elaborate analysis therefore highlights unique trigger points in a live cell that initiate p35 release. We selected two representative drugs that consistently promoted the greatest fold change in p35 release compared to untreated controls, CB-5083 and ionomycin, for further investigation into the mechanisms by which they promote p40-independent p35 release.

**Figure 3.**
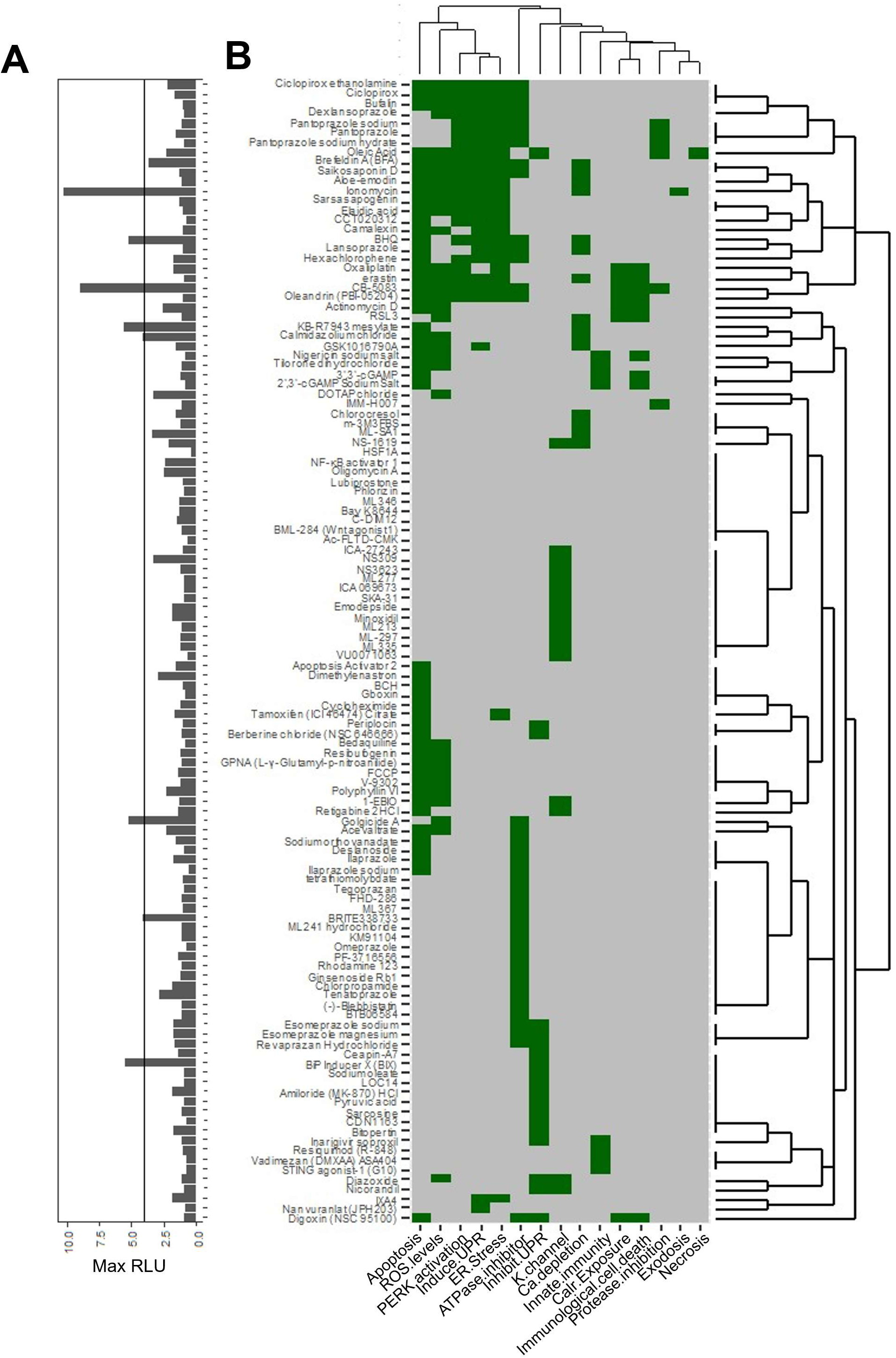
A pharmacological screen reveals key cellular pathways regulating p40-independent p35 release: igh-throughput screening of drugs that could trigger p35 release from L cells. p35NL L cells were plated using Opentrons 2 pipetting robot and exposed to dilutions (100-0.1µm) of 115 drugs for 24 hours. Supernatants were collected after 24 hours of drug exposure for luciferase assay detection of p35NL release (n = 2). Relative luminescence in supernatant of drug-treated p35NL cells was normalized to cells treated with media only. A cut-off for positive inducers of p35 release was set as a 4-fold change and above relative to media only control. Ionomycin was included in the drug screen in addition to its discovery of cell death drugs. **(A)** Maximum fold change from all dilutions (100-0.1µm) of 115 drug library. We identified 8 potential inducers of p35 release (including ionomycin from Figure 2). Line on RLU plot indicates the cut-off of a maximum fold change of greater than 4 relative to a media-only control. **(B)** Unbiased hierarchical clustering of drug functions used in the screen based on available literature.

### Viable cells that do not express p40 can be triggered to release the p35 subunit

In Figures 2 and 3, most drugs screened did not induce significant release of p35 from cells; however, treatment with ionomycin and CB-5083 induced high levels of p35NL release. Ionomycin is known to be calcium ionophore that disrupts intracellular calcium levels, and CB-5083 is an inhibitor of the ER-associated ATPase Valosin-Containing Protein (VCP) (*21*). We validated ionomycin and CB-5083 as inducers of p35NL release and compared their dynamics with NLO release after 24 hours of treatment. p35NL release significantly increased and peaked at 3.12μM ionomycin treatment (Figure 4A, p < 0.0001) with no significant changes in cell viability at this dose (Figure 4B, p = 0.6084). In comparison, L cells only released NLO protein at the highest doses of ionomycin used (50μM) (Figure 4C, p < 0.0001) when viability was completely lost (Figure 4D, p < 0.0001). At the corresponding doses where p35NL viability was lost, there were no significant changes in p35NL release (Figure 4A). These results suggest that ionomycin causes the release of p35NL by an active mechanism in viable cells while NLO is effectively escaping a dying cell passively.

**Figure 4.**
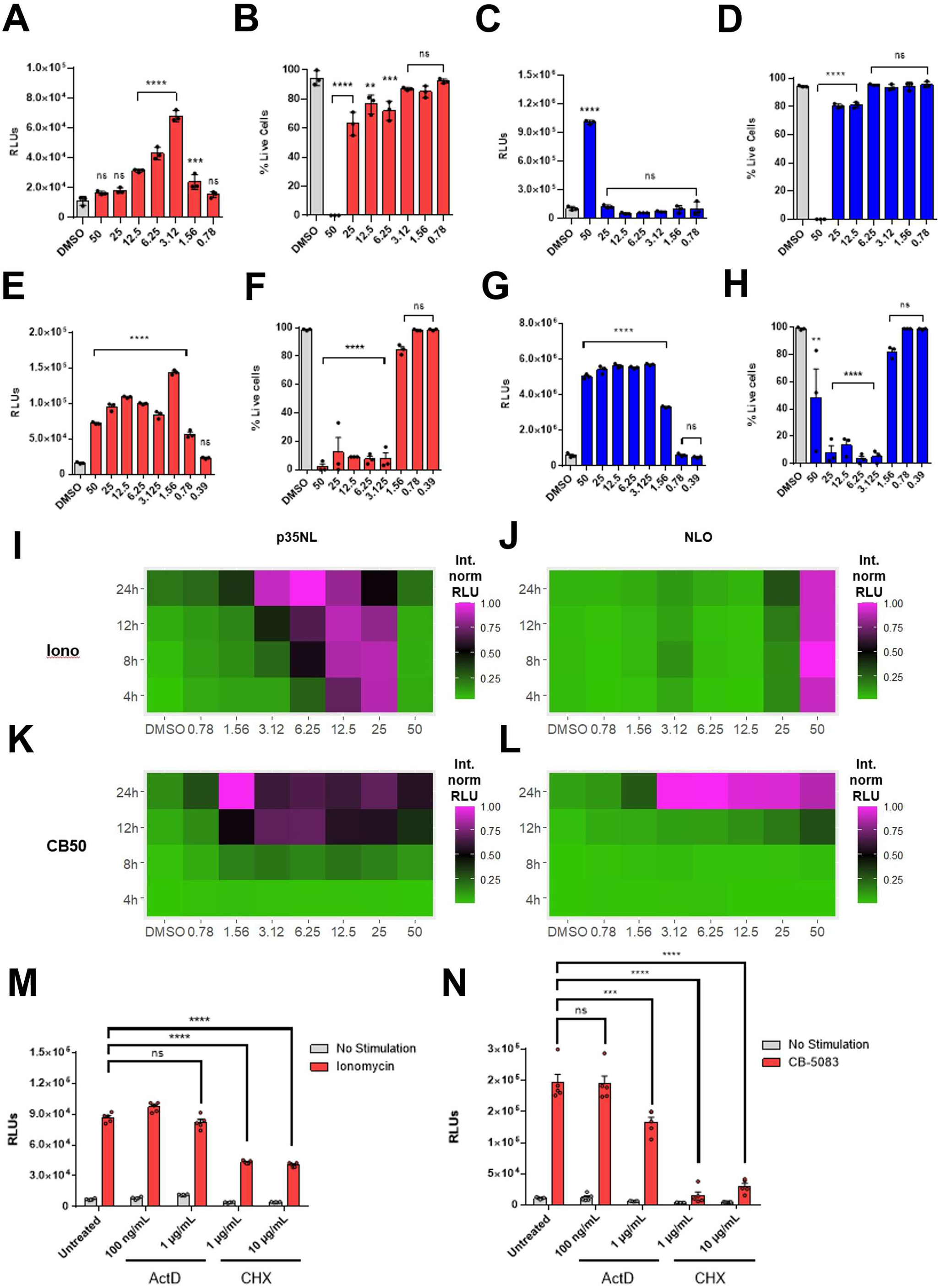
Viable cells that do not express p40 can be triggered to release the p35 subunit: Ionomycin, a calcium ionophore, was validated as a pharmacological trigger of p35 release from viable cells. p35NL and NLO L cells were treated with concentrations of ionomycin ranging from 50-0.78µM. **(A)** Luciferase assay of supernatants from p35NL cells treated with ionomycin for 24 hours **(B)** Viability of p35NL cells after 24-hour ionomycin treatment. **(C)** Luciferase assay of supernatants from NLO cells treated with ionomycin for 24 hours **(D)** Viability of NLO cells after 24-hour ionomycin treatment (n=3). CB-5083, an inhibitor of the multi-functional ATPase VCP (p97), was validated as a pharmacological trigger of p35 release from viable cells. p35NL and NLO L cells were treated with concentrations of CB-5083 ranging from 50-0.39µM. **(E)** Luciferase assay of supernatants from p35NL cells treated with CB-5083 for 24 hours **(F)** Viability of p35NL cells after 24-hour CB-5083 treatment. **(G)** Luciferase assay of supernatants from NLO cells treated with CB-5083 for 24 hours **(H)** Viability of NLO cells after 24-hour CB-5083 treatment (n=3). **(I)** Luciferase assay showing the kinetics of p35NL release in response to ionomycin. **(J)** Luciferase assay showing the kinetics of NLO release in response to ionomycin. **(K)** Luciferase assay showing the kinetics of p35NL release in response to CB-5083 **(L)** Luciferase assay showing the kinetics of NLO release in response to CB-5083 (n=3). RLUs are normalized internally and represented as a heat map **(M)** p35NL cells were pre-treated with media, actinomycin D, or cycloheximide for 60 minutes before the addition of vehicle or 6.25µM ionomycin. Supernatants were collected after 24 hours and used for luciferase assay. **(N)** p35NL cells were pre-treated with media, actinomycin D, or cycloheximide for 60 minutes before the addition of vehicle or 1.56µM CB-5083. Supernatants were collected after 24 hours and used for luciferase assay. Data (A-H, M, N) is represented as raw Relative Luminescence Units. Significant elevations in luminescence were determined through one-way ANOVA of the mean in comparison to vehicle controls (DMSO). Significant changes in cell viability were determined through an ordinary one-way ANOVA of the mean in comparison to vehicle controls (DMSO). *, p < 0.05; ** p < 0.01; *** p < 0.001; **** p < 0.0001; ns = not significant

We investigated the patterns of p35NL release in response to inhibition of VCP with CB-5083. In contrast to ionomycin, CB-5083 promoted p35NL release in a dose-dependent manner both at cytotoxic (50-3.12μM) and viable doses. Indeed, the peak of p35 release was at the dose in which viability was unaffected (1.56μM CB-5083) suggesting that CB-5083 could potentially induce both passive and active modes of p35 release depending on the dose used (Figure 4E, F). NLO cells passively released protein at doses where cell viability was lost (Figure 4G, 4H, p < 0.001).

We then compared the kinetics of p35NL in response to titrations of both ionomycin and CB-5083 treatment at 4, 8, 12, and 24-hr time points. p35NL release began as early as 4 hours after drug treatment with high doses of ionomycin (Figure 4I). Cell viability was decreased in both p35NL and NLO cells in response to the highest dose of ionomycin within 4 hours of treatment (Supplemental Figure 3E, 3F, DMSO vs. 50μM, p < 0.0001 for both p35NL and NLO cell lines). p35 release following CB-5083 treatment was slower and no significant extracellular increase was observed until the 8hr timepoint (Figure 4K). A small but significant elevation in luminescence was observed at the 4-hour timepoints, but it did not increase 3-4-fold until 8 hours (Supplemental Figure 3C). CB-5083 induced slower kinetics of cell death in comparison to ionomycin, as the significant decreases in cell viability did not occur until the 12hr timepoint (Supplemental Figure 3G, DMSO 12hr 50μM, p = 0.0006). The contrast between the release of p35 and NLO are illustrated by comparing Figure 4I vs 4J (for ionomycin) and Figure 4K vs 4L (for CB5083). As indicated by the normalized RLU levels, the release of p35 in the case of ionomycin follows a distinct, almost exclusive pattern in terms of dose and time requirements. The pattern does have some overlap in the case of CB5083; but the peak is still distinct and exclusionary (1.56uM and 24h, see Figure 4K). Together, these results define the timeline of p35 release from these agents - with ionomycin including rapid release within 4 hours, and CB-5083 induced p35 release requiring 8 hours of exposure and leading to cell death within 12 hours of exposure.

The sequential appearance of p35 release and cell death raised the important question of whether (as in the original hypothesis) the former was a consequence of catabolic processes in a cell that was committed to a death program. In other words, is the release of p35 like a stimulus induced secretion of a protein (e.g. cytokine release after PRR signaling) or an early alarmin release from a dying cell. To separate these possibilities, we evaluated the fate of the cells after the inducing compounds (Ionomycin or CB-5083) were washed out of the media. After 8 hours of exposure to ionomycin, removal of ionomycin did not lead to extended release of p35NL indicating that majority of p35NL was released in the first 8 hours of drug exposure and washed off compared to the full 24-hour exposure (Supplemental Figure 4A). Additionally, cell viability could not be rescued in either p35NL or NLO cells after the removal of 50µM ionomycin indicating that the cell death program was already fully initiated at this high dose and could not be rescued past this point (Supplemental Figure 4C, 4D). After 8 hours of exposure of CB-5083 and removal, both p35NL and NLO protein followed similar patterns of release. p35NL release continued after removal of CB-5083, but only at high doses of CB-5083 where viability was still decreased after 8 hours of exposure and then removal (50-12.5µM). This pattern was also seen in NLO protein release where NLO protein released continued after the removal of CB-5083 but was lost at the doses in which cells “recovered” viability (Supplemental Figure 4E, 4F). Overall, these results suggest that ionomycin induces rapid release of p35NL without the need for cell death. In contrast, CB-5083 treatment may present with two modes of release, where some p35 protein is released passively under the cell death program, while a second more active mechanism allows for p35 release from cells exposed to viable doses of VCP inhibition.

The difference in the kinetics of CB-5083 vs. ionomycin prompted us to examine if de-novo gene expression is required to promote the release of p35. p35NL cells were pre-treated with doses of actinomycin D and cycloheximide to inhibit transcription and translation respectively and then were stimulated with ionomycin and CB-5083. In response to ionomycin, inhibition of transcription did not affect the amount of p35 that was released, indicating that active transcription was not necessary to promote p35 release in response to ionomycin stimulation. Pre-treatment with cycloheximide did significantly reduce p35 release during ionomycin-induced release; however, it was not completely abolished (Figure 4M). In sharp contrast, pre-treatment actinomycin D did significantly reduce p35 release in response to CB-5083 at the higher dose. Additionally, inhibition with cycloheximide resulted in a dramatic reduction in p35 release (Figure 4N). Together, these results indicate that VCP inhibition appears to rely on active gene expression to promote p35 release while ionomycin-induced release does not.

### Ionomycin and VCP inhibition induce specific “secretomes” in addition to p35 release

Based on the observations that both ionomycin and CB-5083 treatment induced p35 release from viable cells, but with different kinetics and gene-expression requirements, we investigated whether ionomycin or CB-5083 treatment resulted in a unique profile of other released proteins, or “secretome”. We profiled the supernatants of p35NL L cells treated with ionomycin or CB-5083 using 2D-gel electrophoresis and compared them to supernatants from untreated p35NL L cells. Overall, ionomycin and CB-5083 shared many released proteins in comparison to untreated cells (Supplemental Figure 5A). Ionomycin treatment resulted in very few unique, released proteins that were not found in CB-5083 treatment (Supplemental Figure 5A, 5B), while CB-5083 had variable, increased proteins found in the supernatant that were not present in ionomycin treatments or controls. Together, these data suggest that both treatments result in the release of other proteins besides p35, but only CB-5083 treatment resulted in a subtly unique “secretome”, perhaps due to its transcriptional impact.

### p40-independent p35 release occurs in absence of UPR signaling

The results from our drug screen implicated two major events, intracellular calcium disruption and VCP inhibition, as signaling nodes regulating p40-independent p35 release. Given that the balance of cellular ions and action of proteins such as VCP are critical for the homeostasis of the ER, it was possible that both perturb generic UPR pathways (although such pathways typically lead to translational arrest and cell death, not increased secretion) during p35 release. In our high-throughput screen, some individual activators of ER stress or UPR were tested; but these (e.g. PERK activator, Drug 10: CCT020312) did not induce p35 release. We developed L cells that are stably transduced with a reporter for activation of the IRE1α-XBP-1 axis of the UPR response (*22*). These cells become GFP+ when XBP-1 binds to the UPRE (Unfolded Protein Response Element). The response of UPR-GFP L cells to a time course treatment of ionomycin and CB-5083 was characterized. Like the kinetics of p35 release, ionomycin and CB-5083 treatment induced the UPR with different kinetics. After 8 hours, ionomycin induced a significant increase in the percentage of UPR+GFP+ L cells in comparison to DMSO-treated controls (Figure 5A, p < 0.0001) while the percentage of UPR+ cells in the CB-5083 treatment remained unchanged (Figure 5A, p = 0.3750). This difference was maintained at the 12-hour timepoint. By 24 hours, ionomycin-treated cells had resolved the XBP-1 response while CB-5083-treated cells showed an increased percentage of UPR+GFP+ cells (Figure 5A, p < 0.0001). Overall, these results suggest that both ionomycin stimulation and VCP inhibition can induce the UPR response with kinetics that correspond to those seen in p35NL release.

**Figure 5.**
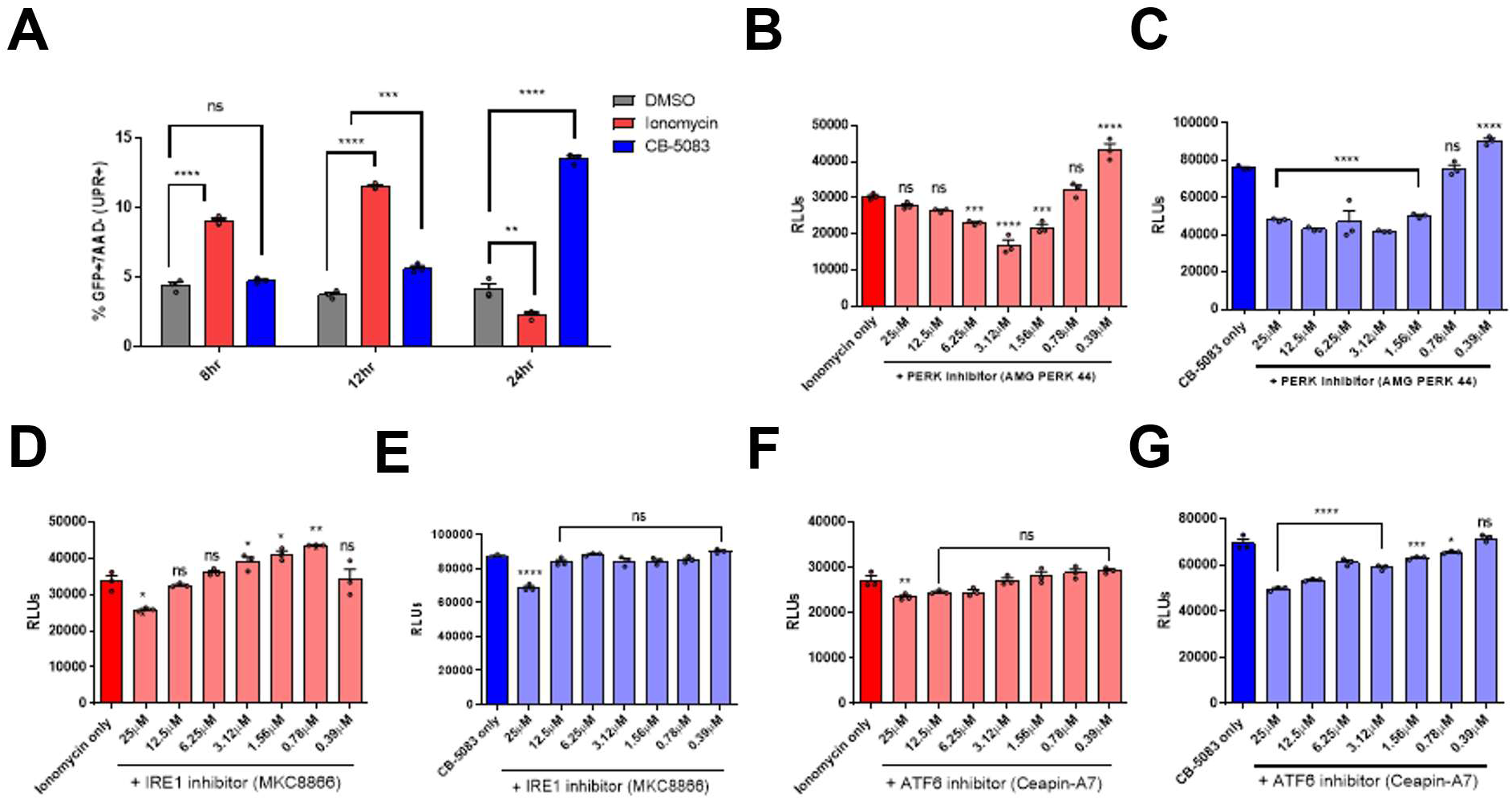
p40-independent p35 release occurs in absence of UPR signaling: (A) Percentage of UPR+ 7-AAD-L cells (GFP+ for XBP-1 binding to Unfolded Protein Response Element) after treatment with vehicle (DMSO), 3.12μM ionomycin, or 1.56μM CB-5083 then analyzed with flow cytometry. (B) Luciferase assay of supernatants from p35NL cells pre-treated with PERK inhibitor AMG PERK 44 for 2 hours then stimulated with 3.12 μM ionomycin for 24 hours. (C) Luciferase assay of supernatants from p35NL cells pre-treated with PERK inhibitor AMG PERK 44 for 2 hours then stimulated with 1.56μM CB-5083 for 24 hours. (D) Luciferase assay of supernatants from p35NL cells pre-treated with IRE1 inhibitor MKC8866 for 2 hours then stimulated with 3.12 μM ionomycin for 24 hours. (E) Luciferase assay of supernatants from p35NL cells pre-treated with IRE1 inhibitor MKC8866 for 2 hours then stimulated with 1.56μM CB-5083 for 24 hours. (F) Luciferase assay of supernatants from p35NL cells pre-treated with ATF6 inhibitor Ceapin-A7 for 2 hours then stimulated with 3.12 μM ionomycin for 24 hours. (G) Luciferase assay of supernatants from p35NL cells pre-treated with ATF6 inhibitor Ceapin-A7 for 2 hours then stimulated with 1.56μM CB-5083 for 24 hours. (n=3). Data is represented as raw Relative Luminescence Units (RLUs). Significant changes in luminescence were determined through ordinary one-way ANOVAs of cells stimulated with ionomycin only or CB-5083 only (control). *, p < 0.05; ** p < 0.01; *** p < 0.001; **** p < 0.0001; ns = not significant

The results that ionomycin and CB-5083 treatment induce the UPR response are not surprising given that previous studies have shown that drugs which induce ER calcium depletion activate the UPR (*23*). Additionally, silencing VCP with shRNA and use of VCP inhibitors has been shown to activate portions of the UPR (*24*). Despite this, it was unclear whether the mechanisms supporting p35 release in response to ionomycin and VCP inhibition require the downstream signaling responses that are propagated by activation of UPR sensors. Disruption of ER homeostasis and activation of UPR is caused by the dissociation of three sensor proteins (ATF6, IRE1α, and PERK) from their chaperone BiP. To determine whether the signaling pathways associated with each sensor in the UPR were essential for promoting independent p35 release, p35NL and NLO L cells were pre-treated with inhibitor for PERK, ATF6, or IRE1α signaling, and then stimulated with 3.12μM ionomycin or 1.56μM CB-5083 for 24 hours. Luciferase assays were performed to determine if the presence of UPR inhibitors decreased p35 release in response to ionomycin or CB-5083.

Signaling through PERK was inhibited through the selective inhibitor AMG PERK 44. First, we confirmed that treatment with AMG PERK 44 alone did not lead to significant amounts of p35 release (Supplemental Figure 6A). Interestingly, PERK inhibition had slightly different effects on p35 release depending on whether cells were treated with ionomycin or CB-5083. During ionomycin stimulation, the highest concentration of PERK inhibitor used (25μM) did not have a significant effect on p35 release (Figure 5B, p = 0.6574), but p35 release was maximally suppressed by ∼50% in response to lower levels of PERK inhibition. In response to CB-5083 treatment, PERK inhibition had a broader, yet less pronounced effect on p35 release. Concentrations of PERK inhibitor ranging from 1.56-25μM suppressed p35 release by ∼30-40%. Regardless, in both situations p40-independent p35 release continued in the presence of PERK inhibition, suggesting that signaling through PERK is not required for p35 release during ionomycin and CB-5083 stimulation.

Next, we tested whether the IRE1α signaling axis was associated with supporting p35 release. p35NL and NLO cells were treated with a selective IRE1α inhibitor (MKC8866) in the presence of ionomycin and CB-5083. Treatment of p35NL and NLO cells with MKC8866 did not influence p35NL or NLO protein release on its own (Supplemental Figure 6K, 6O). Compared to PERK, inhibition of IRE1α appeared to have little effect on p35 release, as in both ionomycin and CB-503 stimulation, significant suppression of p35 release was only seen at the highest dose of inhibitor used (Figure 5D, 5E), a dose at which proliferation and other cellular functions outside of UPR appear to be affected (*25*). These results suggest that the signaling downstream of IRE1α do not contribute to the mechanisms supporting p35 release during ionomycin stimulation and VCP inhibition.

Finally, the contribution of the ATF6 arm of the UPR to p35 release was tested. p35NL and NLO cells were treated with an ATF6 inhibitor, Ceapin-A7, in the presence of ionomycin and CB-5083. During ionomycin stimulation, ATF6 inhibition caused a small, but significant decrease in the amount to p35NL being released at the highest dose of inhibitor used (Figure 5F). ATF6 inhibition had a more pronounced effect on p35 release during CB-5083 treatment, as suppression of p35 release was seen at doses ranging from 0.78-25μM Ceapin-A7 (Figure 5G). Like PERK inhibition, ATF6 inhibition during CB-5083 stimulation led to variable decreases in p35NL and NLO cell viability (Supplemental Figure 6Y, Z) suggesting that at least some of the decrease in p35NL release observed can be attributed to the death of cells rather than an active mechanism of suppression. Together, the sum of these experiments suggests that while reduction can be observed in response to inhibition of the individual arms of the UPR, p35NL release continues without a strict reliance of UPR pathways.

### Ionomycin stimulation induces aggregates of p35-Scarlet around Golgi markers

Given that ionomycin and CB-5083 induced release of p35 from viable cells, more analogous to cytokine secretion than a death-associated failure of intracellular retention, we proceeded to map the intracellular trajectory of p35 in response to these drugs using the p35-Scarlet reporter cells described in Figure 1. Cells were exposed to 3.12μM ionomycin (the dose at which we observed the highest levels of p35 release) for 24 hours, and the localization of p35-Scarlet was tracked. In response to ionomycin stimulation, we observed aggregation of p35-Scarlet in a one aggregate rather than distributed throughout both poles of the cell like in DMSO-treated controls. The aggregates of p35-Scarlet appeared to localize around corresponding Golgi-eGFP markers despite some puncta of p35 occurring outside of the Golgi markers (Figure 6A).

**Figure 6.**
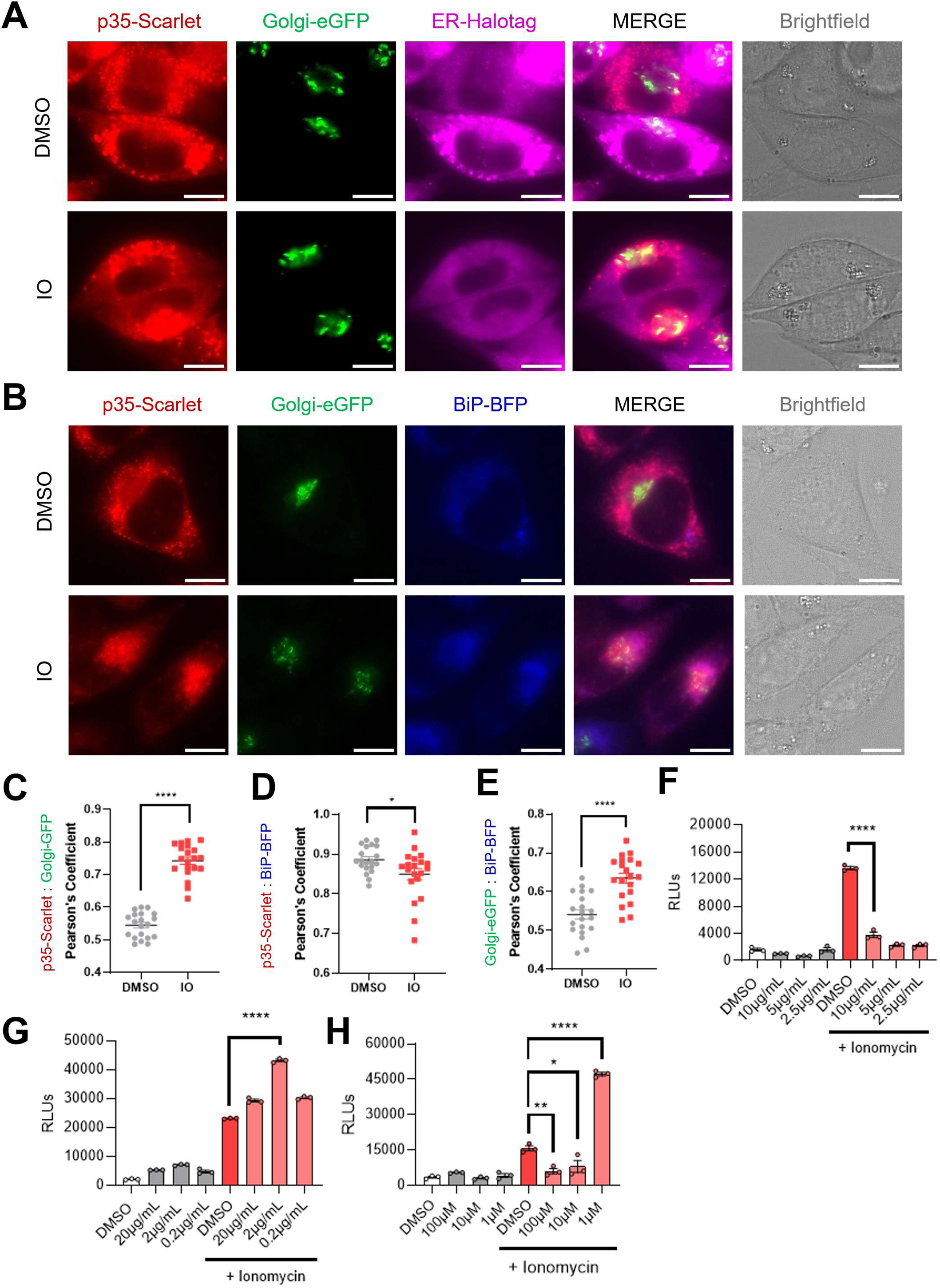
Ionomycin stimulation induces aggregates of p35-Scarlet around Golgi markers: Subcellular localization of p35 under ionomycin-induced release conditions. **(A)** Representative live cell imaging of p35-Scarlet/Golgi-eGFP/ER-Halotag L cells treated with either vehicle (DMSO) or 3.12μM ionomycin (IO) for 24 hours. 10μm scale bars. **(B)** Representative live cell imaging of p35-Scarlet/Golgi-eGFP/BiP-BFP L cells treated with treated with either vehicle or 3.12μM ionomycin for 8 hours. 10 μm scale bars (n=20). **(C)** Quantification of colocalization between p35-Scarlet and Golgi-eGFP using Pearson’s coefficient (n=20). **(D)** Quantification of colocalization between p35-Scarlet and BiP-BFP using Pearson’s coefficient (n=20). **(E)** Quantification of colocalization between BiP-BFP and Golgi-eGFP using Pearson’s coefficient (n=20). **(F)** Luciferase assay of p35NL supernatants from cells pre-treated with COP-I inhibitor (Brefeldin A) for 2 hours and then stimulated with 6.25μM ionomycin for 4 hours (n=3). **(G)** Luciferase assay of p35NL supernatants from cells pre-treated with NFAT inhibitor, cyclosporin A (CsA) for 2 hours and stimulated with 6.25μM ionomycin for 4 hours (n=3). **(H)** Luciferase assay of p35NL supernatants from cells pre-treated with BAPTA-AM for 30 minutes and stimulated with 6.25μM ionomycin for 6 hours. (n = 3). Significant changes in colocalization were determined using an unpaired t-test of the mean in comparison to vehicle controls (DMSO). Each image analyzed was a 100X frame containing 5-10 cells. *, p < 0.05; ** p < 0.01; *** p < 0.001; **** p < 0.0001; ns = not significant

Both p35 and p40 subunits interact with several ER chaperones during their coordinated folding and linkage. It has been hypothesized that association with ER chaperones is the primary restrictive mechanism that prevents the independent release of p35 in the absence of p40, although which ER chaperone plays the largest role in its retention is not clear (*14*). Therefore, we hypothesized that interactions with important ER chaperones such as BiP may change during independent release of p35 in response to stimulation with ionomycin. Once again, ionomycin induced the accumulation of p35-Scarlet near the Golgi-eGFP marker protein as previously shown (Figure 6B). This was quantified by a significant increase in Pearson’s correlation coefficient in ionomycin-treated vs vehicle L cells between p35-Scarlet and Golgi-eGFP (Figure 6C, p < 0.0001). After 8 hours of stimulation with ionomycin, BiP-BFP trafficked similarly to p35-Scarlet where it formed a single aggregate that appeared to overlap with p35-Scarlet and Golgi-eGFP (Figure 6B). There was a slight but significant decrease in the colocalization between p35-Scarlet and BiP-BFP (Figure 6D, p = 0.0264). When comparing the overlap between BiP-BFP and Golgi-eGFP using Pearson’s coefficient of correlation, we observed that ionomycin induced a significant increase in the correlation between BiP-BFP and Golgi-eGFP (Figure 6E, p < 0.0001). Together, these results suggest that both p35-Scarlet and BiP-BFP may traffic to the Golgi during stimulation with ionomycin.

Based on the data suggesting that p35-Scarlet may traffic through the Golgi during release in response to ionomycin, we further characterized the mechanisms required for ionomycin-induced release. p35NL cells were pre-treated with the COP-I inhibitor, Brefeldin A, and then stimulated with ionomycin to determine whether COP-I dependent trafficking is required for p35NL release in response to ionomycin. Stimulation with ionomycin led to p35NL release, but this was significantly suppressed in the presence of BFA pre-treatment (Figure 6F, p < 0.0001). These results indicate that retrograde transport may be necessary for p35 release in response to ionomycin.

The intracellular calcium flux generated by stimulation with ionomycin can initiate several calcium-dependent signaling pathways. One such pathway involves the activation of the transcription factor, NFAT, which is activated by signaling through calmodulin-calcineurin complexes after intracellular calcium increase. We determined whether calcineurin/NFAT signaling activated by drugs like ionomycin was necessary for p35 protein to be released when cells are stimulated with ionomycin. p35NL cells were pre-treated with the calcineurin inhibitor, Cyclosporin A, and then stimulated with ionomycin. Surprisingly, inhibition of NFAT signaling did not suppress p35NL release in response to ionomycin; instead, we observed a significant increase in release when 2µg/mL CsA was combined with ionomycin in comparison to ionomycin alone (Figure 6G, p < 0.0001).

Additionally, we observed the response of p35 release during stimulation with ionomycin in the presence of a calcium chelator, BAPTA-AM. Pre-treatment with higher concentrations of BAPTA-AM led to significantly decreased release of p35NL during ionomycin stimulation. Surprisingly, the lowest concentration of BAPTA-AM used induced the opposite effect where presence of both BAPTA-AM (1μM) led to approximately a 3-fold increase in p35 release compared to ionomycin stimulation alone (Figure 6H).

Together, these results suggest that stimulation with ionomycin induces the translocation of p35-Scarlet from its ER localization to aggregation with Golgi markers. This translocation appears to be COP-I dependent yet independent of calcineurin-associated signaling pathways. Chelation of calcium with specific doses of BAPTA-AM reduced p35 release in response to ionomycin suggesting that intracellular calcium flux was critical for the release mechanism.

### VCP inhibition leads to aggregation of p35-Scarlet near Golgi markers and the formation of unique vacuolated structures

Next, we characterized the trajectory of p35-Scarlet in response to treatment with the VCP inhibitor, CB-5083. p35-Scarlet reporter cell lines were exposed to 1.56µM CB-5083 for 24 hours and were observed using live-cell fluorescent imaging. Similar to ionomycin, treatment with CB-5083 led to aggregates of p35-Scarlet that appeared to localize near Golgi-eGFP (Figure 7A).

**Figure 7.**
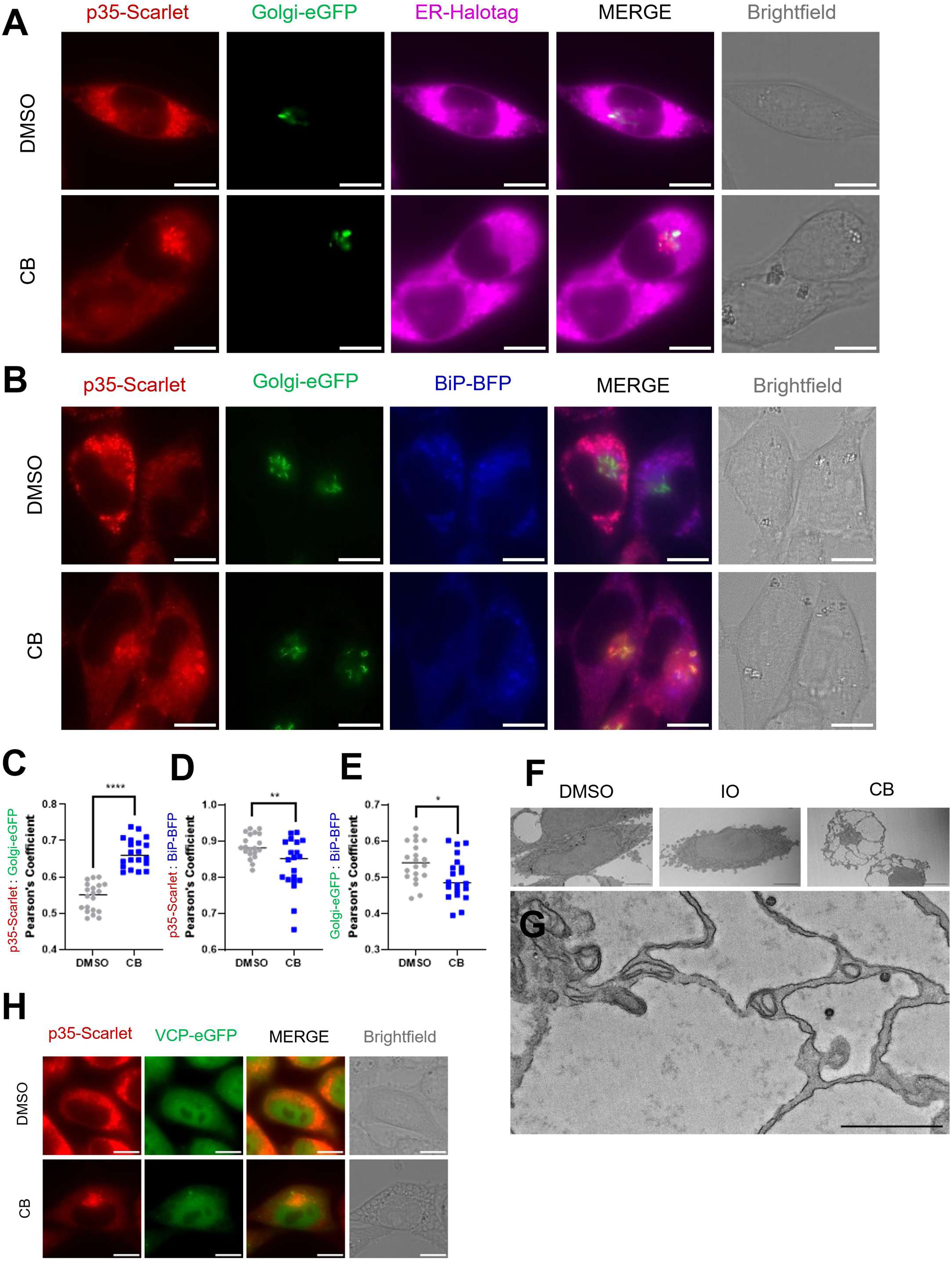
VCP inhibition leads to aggregation of p35-Scarlet near Golgi markers and the formation of unique vacuolated structures. Subcellular localization of p35 under CB-5083-induced release conditions. **(A)** Representative live cell imaging of p35-Scarlet/Golgi-eGFP/ER-Halotag L cells treated with either vehicle (DMSO) or 1.56μM CB-5083 (CB) for 24 hours. 10 μm scale bars **(B)** Representative live cell imaging of p35-Scarlet/Golgi-eGFP/BiP-BFP L cell treated with treated with either vehicle or 1.56μM CB-5083 for 8 hours (n=20). 10 μm scale bars **(C)** Quantification of colocalization between p35-Scarlet and Golgi-eGFP using Pearson’s coefficient (n=20). **(D)** Quantification of colocalization between p35-Scarlet and BiP-BFP using Pearson’s coefficient (n=20). **(E)** Quantification of colocalization between BiP-BFP and Golgi-eGFP using Pearson’s coefficient (n=20). **(F)** Transmission electron microscopy of L cells treated with vehicle, 3.12μM ionomycin, or 1.56μM CB-5083 for 24 hours. **(G)** Ultrastructure of vacuoles from CB-5083-treated L cells after 24 hours. **(H)** Representative live cell images from p35-Scarlet/VCP1-eGFP L cells treated with either vehicle, 3.12μM ionomycin, or 1.56μM CB-5083 for 12 hours (n=1). Significant changes in colocalization were determined using an unpaired t-test of the mean in comparison to vehicle controls (DMSO). Each image analyzed was a 100X frame containing 5-10 cells. *, p < 0.05; ** p < 0.01; *** p < 0.001; **** p < 0.0001; ns = not significant

We compared the interactions of p35-Scarlet with its ER chaperone BiP under CB-5083 release conditions. CB-5083 treatment increased the localization of p35-Scarlet with Golgi-eGFP (Figure 7B, 7C). In contrast to ionomycin results, correlation between BiP-BFP and Golgi-eGFP did not increase in response to CB-5083 (Figure 7E); in fact, it was significantly decreased in comparison to DMSO-treated controls. Overall, these results could suggest several things: 1) ionomycin promotes the release of both BiP-GFP and p35-Scarlet through conventional secretory through Golgi-mediated mechanisms 2) VCP inhibition does not promote the same mechanism leading to the release of p35 since its interactions are different with BiP upon release 3) Kinetics of BiP release may be slower under VCP-inhibiting conditions, given that we saw slower kinetics of cell death and p35 release in CB-5083 treated cells in comparison to ionomycin.

CB-5083 is a phase I chemotherapy candidate, due to its cytotoxic effects on cancer cells. However, we observe functional p35 release at non-cytotoxic doses. VCP performs several homeostatic roles in the cell, and it has already been shown to interact with similar IL-12 family subunits (*26*). Therefore, we investigated how the VCP protein itself may interact with p35 at baseline and in the context of independent release conditions. We generated L cells that stably express fusion proteins p35-Scarlet and VCP-eGFP. We compared the localization of p35-Scarlet and VCP-eGFP under treatment with 1.56uM CB-5083 after 12 hours. In vehicle-treated cells, VCP was distributed widely throughout the cell with specific puncta in the cytoplasm; however, it did not appear to directly localize with p35-Scarlet. Similar to the reporter experiments, p35-Scarlet localized into a central aggregate after 12 hours of treatment with CB-5083. Specifically in the CB-5083 treated group, VCP inhibition induced the formation of vacuolated structures throughout the cell that were visible in Brightfield (designated by yellow arrow) (Figure 7H). These structures were not present in the doses of ionomycin that we examined; however, we cannot rule out the possibility that higher doses of calcium disruption may induce them. We investigated how the trajectory of these structures corresponds with p35 release. L cells were treated with DMSO, ionomycin, and CB-5083 for 24 hours and then sectioned for thin layer transmission electron microscopy. At this resolution of electron micrographs, we did not observe similar structures in ionomycin-treated cells, at doses associated with p35 release. The structures formed after CB-5083 treatment appeared as large, vacuolated structures that were bordered by electron dense membranes but lacked significant density inside, suggesting that they are perhaps fluid rich but devoid of dense protein or lipid structures (Figure 7G). Together our data suggests that in addition to a conventional secretion (via the Golgi) VCP inhibition induces the formation of membranous, vacuole-like structures. Currently there are no reagents we are aware of to directly probe whether the latter vesicles are necessary for p35 release.

### The mechanisms of p35 release identified here operate in primary murine tissues

The systematic pharmacogenomic approaches we have used so far allowed us to identify pathways that regulate p35 release using murine L cells, which are derived and immortalized from adipose tissue. To extend these studies into primary tissues, we generated primary cells from mice. Muscle fibroblasts were derived from postnatal day 3 p40KO mice (to ensure that the release was p40-independent) and transduced with either p35NL or NLO retrovirus virus to express the reporter constructs. p35 release in response to a titration of ionomycin after 24 hours was observed. Similar to L cells, primary p40 KO cells released p35NL protein at doses of drug where there were no significant differences in cell viability (Figure 8A, 8B). Intriguingly though the precise dose of ionomycin that elicited maximal release was lower than it was L cells (perhaps indicating a differential sensitivity of primary muscle cells), the pattern and sharp threshold effects were both replicated in primary cells (Figure 7A). NLO protein was still released at lethal doses of ionomycin (Figure 8C, 8D).

**Figure 8.**
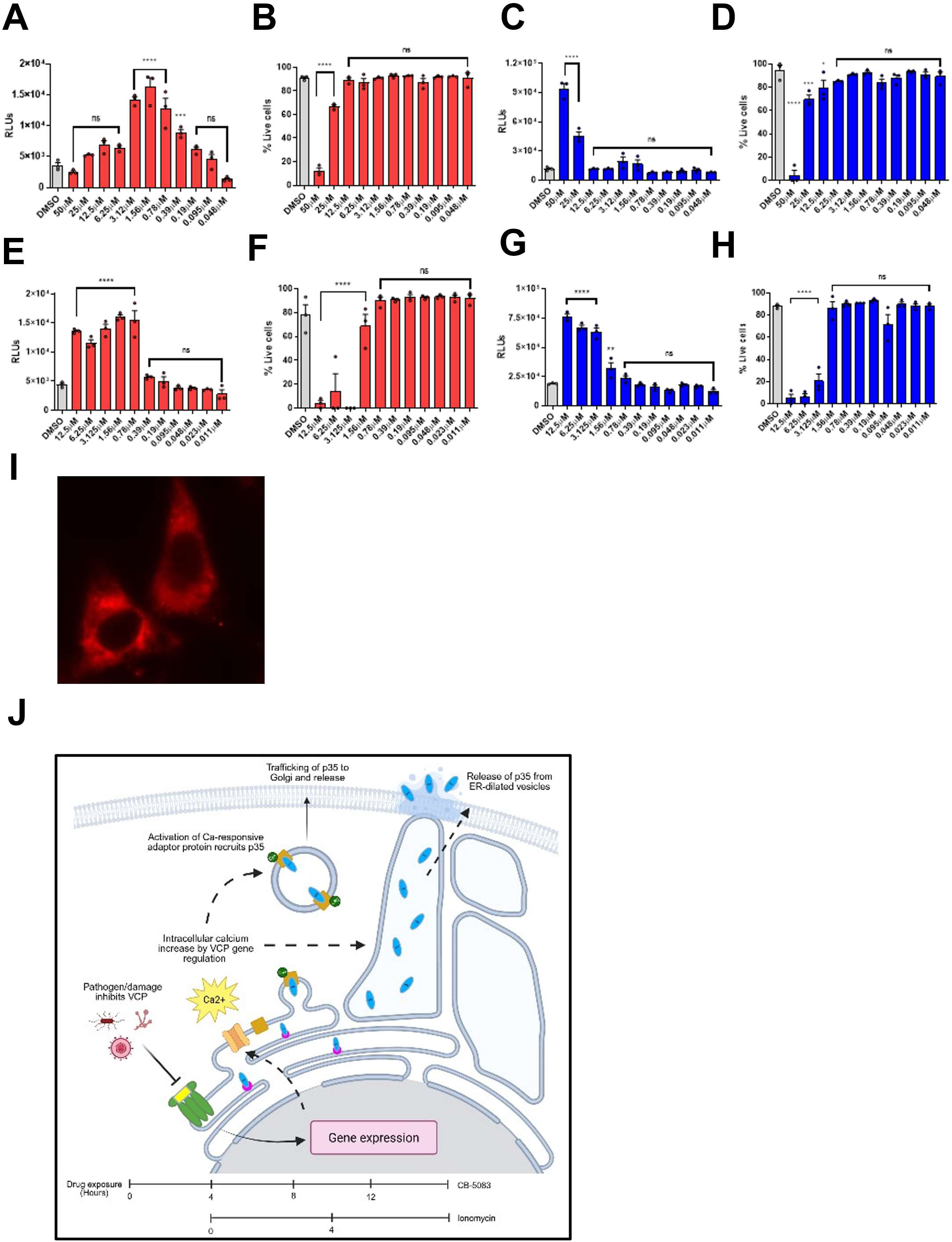
The mechanisms of p35 release identified here operate in primary murine tissues. **(A)** Luciferase assay of supernatants from p40KO muscle fibroblasts transduced with p35NL and treated with titration of ionomycin for 24 hours. **(B)** Viability of p35NL primary p40KO fibroblasts treated with ionomycin for 24 hours. **(C)** Luciferase assay of supernatants from p40KO muscle fibroblasts transduced with NLO and treated with titration of ionomycin for 24 hours. **(D)** Viability of NLO primary p40KO fibroblasts treated with ionomycin for 24 hours. **(E)** Luciferase assay of supernatants from p40KO muscle fibroblasts transduced with p35NL and treated with titration of CB-5083 for 24 hours. **(F)** Viability of p35NL primary p40KO fibroblasts treated with CB-5083 for 24 hours. **(G)** Luciferase assay of supernatants from p40KO muscle fibroblasts transduced with NLO and treated with titration of CB-5083 for 24 hours. **(H)** Viability of NLO primary p40KO fibroblasts treated with CB-5083 for 24 hours. **(I)** Live cell imaging of p40KO muscle fibroblast transduced with p35-Scarlet and imaged at 100X magnification. **(J)** A theoretical model for independent p35 release initiated by pathogen-induced VCP inhibition. We propose a model of p35 release from stromal cells which is initiated by pathogen-induced VCP inhibition. Inhibition of VCP promotes gene expression that increases intracellular calcium flux, potentially through the upregulation of an ER-associated channel like IP_3_R. From this point, our data suggests two possible modes of release. In the first, intracellular calcium activates a calcium-responsive cargo adaptor and promotes the selective passage of p35 through the conventional secretory pathway. In a second mode of release, we propose that the vacuole-like structures induced by VCP inhibition may be critical in allowing a Golgi-independent pathway to release of p35. In this mechanism, an imbalance of intracellular calcium promoted by VCP inhibition leads to the dilation of the ER into vacuoles containing p35. ER dilation promotes the release of p35 through an unknown, perhaps novel pathway in manner that does not rely on cell death. Data is represented as raw Relative Luminescence Units. Significant elevations in luminescence were determined through one-way ANOVA of the mean in comparison to vehicle controls (DMSO). Significant changes in cell viability were determined through an ordinary one-way ANOVA of the mean in comparison to vehicle controls (DMSO). *, p < 0.05; ** p < 0.01; *** p < 0.001; **** p < 0.0001; ns = not significant

In response to CB-5083 treatment, primary p40 KO cells released p35NL protein both at doses that resulted in cell death and at doses where viability was restored (Figure 8E, 8F). Primary cells were also more sensitive to VCP inhibition as the release of p35NL peaked at 0.78μM rather than 1.56μM. NLO protein was released doses where cell viability was lost (Figure 8G, 8H). Finally, we also validate the localization studies using primary p40KO cells transduced with p35-Scarlet retrovirus. Live-cell imaging revealed that like reporter cell lines, p35-Scarlet protein was distributed and retained throughout the cytosol (Figure 8I).

Taken together these data offer insights into the intracellular pathways regulating the retention and release of p35 in non-immune stromal cells (Figure 8J) – both in vitro as well as ex-vivo primary tissues.

## DISCUSSION

The importance of the IL-12 family as key “bridge cytokines” connecting early innate responses with the differentiation of adaptive cells has spurred significant interest in understanding the triggers that lead to their production. The secretion of the p40 subunit follows a conventional pathway downstream of PRRs and NFκB-dependent transcription (*27*). In contrast, understanding how the corresponding p35 subunit is sourced has not been straightforward. Despite initial reports that DCs secrete bioactive IL-12 heterodimers after PRR signaling, we and others have shown that this requires additional signals such as CD40L and IFN-γ (*28, 29*). Furthermore, the identification of a separate pathway (two-cell IL-12) whereby stromal cells contribute p35 to complement haematopoietically-derived p40 exposed the fundamental lack of clarity on the mechanisms operating in stromal cells to allow for p35 release (*4, 15*). A particular challenge in this regard is that in the absence of p40, IL-12p35 is retained in the ER, misfolded, and thought to be subject to degradation by proteins in the ERAD pathway (*10, 26*). Since the ER regulates an extensive catalog of cellular processes ranging from stress sensing, metabolic control, UPR, and even initiates various forms of cell death, pinpointing a precise mechanism that allows for p35 release is not trivial. In this context, the new model we propose involving VCP activity and intracellular ionic flux (Figure 8J) in driving active p35 secretion from live stromal cells is foundational for expanding our understanding of the cellular mechanisms regulating IL-12 production.

The ability of ionomycin, a calcium ionophore (*30, 31*), to trigger p35 release (Figures 2, 4), theoretically seemed consistent with a mechanism that disrupted the relationship of p35 with ER chaperones such as BiP, releasing it from ER retention and allowing trafficking to the Golgi for secretion (*32–34*). Indeed, BiP is a central player that associates with UPR sensors and triggers the activation of that pathway and even cell death in response to metabolic stress (*35–37*). While many of these pathways appear interconnected, each also has distinct physiological roles (*38*). Intriguingly, the calcium-dependent release of p35 (inhibited by high concentrations of calcium chelator BAPTA-AM, Fig. 6H) seems to be independent of calcium’s role in triggering UPR or cell death (Figures 4, 5). Previous studies have demonstrated the role of calcium in promoting the dissociation of BiP from ER-restricted “clients” which results in the secretion of proteins like the TCRα chain. In these studies, decreasing intracellular calcium (by chelation with BAPTA-AM) was sufficient to promote secretion of client proteins. However, BAPTA-AM by itself does not induce the release of p35, suggesting that the significance of calcium in promoting release of p35 is perhaps compartmentalized. It is also not dependent on Ca^2+^ flux driven gene expression, as blocking the NFAT pathway (Figure 6F) or just global transcription (Figure 4M) was not relevant. Perhaps this is indicative of the involvement of an as-yet-unidentified cytosolic or ER outer-membrane calcium-responsive adaptor (as shown in Figure 8J). Candidates for such an adaptor include ALG-2 which has been shown to dynamically adjust transport from the ER to Golgi based on intracellular calcium flux (*39*) but require further studies. Importantly, both the sharp triggering dose of ionomycin as well as the counter-intuitive increase in p35 release at low doses of BAPTA-AM (1μM) in the presence of ionomycin suggests that the proposed adaptor mechanism has a narrow threshold for activation.

Staging the role of VCP (p97) – an ATPase with multiple roles in cellular morphology, organelle biology, and gene expression (*21, 40, 41*) is facilitated by our finding that VCP inhibition leads to maximal p35 release in a gene-expression dependent manner (Figure 4N). Cells subject to treatment with CB-5083 also display altered morphology with exaggerated, vacuolated structures (Figure 7F). An important question, albeit one challenging to resolve at this stage, is whether the targets of VCP inhibition and ionomycin operate in the same linear pathway that promotes p35 release. Based on the differences in the ability of cycloheximide and actinomycin D treatment to suppress p35 release in CB-5083-treated cells with decreased effect on ionomycin-induced release (Figure 4M, N), one could envision that VCP inhibition promotes the expression of either a regulator of the cytosolic calcium adaptor or an alternate channel that translocates intracellular calcium to the appropriate compartment without ionomycin (Figure 8J).

Overall, the combined data from CB-5083 and ionomycin treatments allows us to form a theoretical framework for the independent release of p35 in the context of two-cell IL-12. In this model, p35 release is initiated by intracellular infection or damaged associated with infection leading to the inhibition of VCP. Inhibition of VCP promotes gene expression that increases intracellular calcium flux, potentially through the upregulation of an ER-associated channel like IP_3_R. From this point, our data suggests two possible modes of release. In the first, intracellular calcium activates a calcium-responsive cargo adaptor and promotes the selective passage of p35 through the conventional secretory pathway. This model is supported by our imaging data showing aggregation of p35 around Golgi markers during drug treatments (Figure 6B and 7B), and the inhibition of p35 release when exposed to the secretion blocker, BFA, though alternate effects of BFA may affect intracellular levels of p35(*42*). In a second mode of release, we propose that the vacuole-like structures induced by VCP inhibition (Figure 7H) may be critical in allowing a Golgi-independent pathway to release of p35. In this mechanism, imbalance of intracellular calcium promoted by VCP inhibition leads to the dilation of the ER into vacuoles containing p35. These structures are consistent with a mechanism of cell death termed, paraptosis, where calcium imbalance leads to the extreme dilation of the ER and mitochondria (*43, 44*). While these structures are typically associated with cell death, we proposed that these dilated vesicles may be associated with a novel trafficking mechanism that promotes p35 release without cell death (Figure 8J).

A clear resolution provided by our data regards the role of active vs passive release of p35 from non-immune cells (Figure 2), as multiple methods of inducing cell death did not lead to large amounts of p35 release compared to NLO controls. This is significant, as prior hypothesis about the release of p35 in the context of two-cell formation of IL-12 hinged on the potential role of cell death associated with infection (*4, 15*). Most significant of all was the observation that induction of cell death using ionomycin did not lead to any release of p35NL in comparison to vehicle controls. There is evidence that ionomycin-induced cell death activates endogenous calcium-activated calpains (*45*), but whether or not these proteins can contribute to the active degradation of p35 during cell death is not clear. Additionally, the same relationship is not observed in lethal doses of CB-5083. Overall, our comparisons of the kinetics of p35 release in response to ionomycin vs. CB-5083 show a distinct negative correlation between cell death and p35NL release where the highest levels of release are associated with restoration of cell viability (Supplemental Figure 3 I, K)

Recent reports also implicate VCP1 in pathogen sensing, signaling of C-type lectin receptors (CLRs) (*46*) or as an antibacterial against intracellular *Salmonella* (*47*). Similarly, *Listeria monocytogenes* (*48*) and *Leishmania donovani* (*49*) infections are reported to cause temporal changes in intracellular calcium homeostasis. Such emergent studies connect with this work, suggesting a precisely calibrated intracellular apparatus that helps non-immune cells instruct the effector functions of local immune cells. Given that CB-5083 and other VCP inhibitors have been heavily investigated for treatment of cancer (*50*), the idea that they may also drive formation of IL-12 in a localized manner is intriguing and warrants further investigation. Canonically, antigen-presenting cells regulate adaptive immunity by bridging affected tissue-level cues with events in SLOs. Damaged tissue cells passively release pro-inflammatory molecules (DAMPs) which activate these DCs. The VCP1/Ca^2+^ flux dependent intracellular sensors allow tissue cells themselves to identify and actively contribute portions of pro-inflammatory cytokines leading to the formation of active IL-12 in tissues, allowing for fine-tuning of effector responses in localized microenvironments.

## MATERIALS AND METHODS

### Mouse models

C57BL/6, SMARTA TCR transgenic (B6.Cg-Ptprca Pepcb Tg(TcrLCMV)1Aox/PpmJ) mice (gift of Dr. Dorian McGavern, NIH Bethesda) were bred to RAG2-KO background as described in (*4*). RAG2-KO (B6.Cg-*Rag2^tm1.1Cgn^*/J) were obtained from Jackson Laboratory and were used as a lymphocyte-free source of splenocytes for antigen presentation in in vitro T cell restimulation assays. IL-12p40-KO mice (B6.129S1-*Il12b^tm1Jm^*/J) were obtained from Jackson Laboratory and used to generate primary muscle cell lines. Mice were housed under IACUC conditions and Specific Pathogen Free (SPF) conditions at the University of Maryland School of Medicine.

### Constructs

The p35-NanoLuc (p35NL) construct was generated by InFusion cloning (Takara) with a GeneBlock (Genscript) containing the murine p35 cDNA linked to NanoLuc luciferase through a serine-glycine linker which was then cloned into the retroviral backbone MIGR1-GFP. The NanoLuc-only (NLO) construct was generated in a similar manner by InFusion cloning of NanoLuc luciferase cDNA only into retroviral backbone MIGR1-GFP. Co-expression of p40 experiments used the murine p40 with a serine-glycine linker cloned into expression vector CMV-eYFP. The p35-Scarlet (p35-Scar) construct was generated by amplifying murine p35 cDNA from a plasmid containing the IL-12 heterodimer cDNA (Addgene #108665) and cloning into MIGR1-GFP with a Myc-mScarlet cDNA fragment amplified from a previous plasmid using InFusion cloning. Imaging reporter cell lines used the following fusion constructs synthesized at Genscript by cloning each fluorescent reporter into MIGR1 retroviral backbones: Golgi-eGFP (sequence targets β-1,4-galactosyltransferase 1), BiP-BFP, ER-Halotag (sequence targets KDEL sequence with HaloTag® from Promega), VCP-eYFP (Genscript). Novel imaging constructs were submitted to Addgene (Catalog numbers). To report on the activation of the unfolded protein response, the construct (lenti-25xUPRE-minP-EGFP) was a gift from Seiichi Oyadomari (Addgene plasmid # 159668; http://n2t.net/addgene:159668; RRID:Addgene 159668) (Kitakaze et al. 2019). Constructs resulting from all InFusion cloning were verified using whole plasmid DNA sequencing from Psomagen.

### Cell Lines

A sub-line of Expi293F™ (referred to as HEK cells), grown in serum (DMEM with 10% FBS) were used for transient expression experiments in Figure 1. Retroviruses were generated using Phoenix-GP cell lines (ATCC - CRL-3215™). Both Expi and Phoenix-GP cells were maintained in DMEM media supplemented with 1% antibiotic/antimycotic, 1% sodium pyruvate, 1% L-glutamine, and 10% fetal bovine serum (Gibco). Murine fibroblast L cells (ATCC - CRL-2648™) were used to generate stable expression of p35-NanoLuc, Nano-Luc only, p35-Scarlet/Golgi-eGFP/ER-Halotag, p35-Scarlet/Golgi-eGFP/BiP-BFP, and UPR-GFP for all imaging and luciferase assays. L cells were maintained in RPMI 1640 supplemented with 1% antibiotic/antimycotic, 1% L-glutamine, and 10% fetal bovine serum (Gibco).

### Transfection of cells

Transient expression of p35-NanoLuc and NanoLuc-only was achieved through transfection of plasmid DNA with polyethyleneimine (PEI) in HEK cells. 1 x 10^5^ HEK cells were plated in 24-well plates (Corning) the day before transfection. Cells were transfected at a ratio of 1:3 plasmid DNA to PEI. For co-expression experiments, HEK cells were transfected with 1μg/mL p40-linker plasmid or empty vector (CMV-eYFP) and increasing concentrations (0.125μg/mL – 2.0μg/mL) of p35-NanoLuc/NanoLuc-only plasmid DNA. Cells were transfected overnight and then media was changed to remove PEI and plasmid DNA. Supernatants for luciferase assay were collected 48 hours after media change. To collect supernatants, plates were spun down at 2000 rpm for 7 min. Supernatants were transferred to microcentrifuge tubes and spun down again at 2000 rpm for 7 min to ensure a cell-free supernatant.

### Retroviral transduction of L cells

All retroviruses used were generated using Phoenix-GP cells. 4 x 10^6^ cells were PEI-transfected with 10μg target plasmid DNA and 20μg of retroviral packaging construct pCL-Eco. Media was changed after overnight transfection, and supernatants containing retrovirus were collected for 3 days and concentrated with RetroX concentrator solution (Takara Bio). L cells were plated in a 24-well plate and transduced with polybrene and retrovirus concentrate at 1:10 ratio. Cells were sorted for stable expression of p35-NanoLuc and NanoLuc-only using the top 5% of GFP+ cells using the Cytek Aurora CS Sorter. Cells used for live-cell, fluorescent imaging were sorted for expression of fluorescent markers proteins using the Cytek Aurora CS Sorter.

### Intracellular cytokine staining (ICS)

An in vitro culture of activated transgenic T cells was generated using lymph nodes collected from a SMARTA transgenic mouse combined with splenocytes from a RAG2-KO mouse. Both lymphocytes and splenocytes were crushed into a single cell suspension using mesh and crushing buffer with 1X PBS, 1% antibiotic-antimycotic, and 5% fetal bovine serum. Lymphocytes and splenocytes were combined in T cell media (RPMI 1640 supplemented with 1% antibiotic/antimycotic, 1% L-glutamine, and 10% fetal bovine serum, and 0.000014% 2-mercaptoethanol), and T cells were activated with 1μM GP_61-80_ antigen. After 3 days of activation, SMARTA T cells were expanded with 10 units/mL of recombinant murine IL-2. 24 hours later, cells were harvested and plated 1 x 10^5^ cells per well in a 96-well round bottom plate (Corning). Cells were restimulated with 1µM GP_61-80,_ 3 x 10^5^ RAG2-KO splenocytes, and either 100ng/mL recombinant p40, 100ng/mL recombinant IL-12, or 50µL of supernatant from HEK cells co-expressing p35-NanoLuc and p40. After 2 hours of restimulation, Brefeldin A was added to stop secretion for 4 hours. Cells were harvested and washed with FACS buffer (1X PBS with EDTA, BSA, and sodium azide). Cells were incubated on ice with FcBlock for 15 minutes to prevent Fc receptor binding before staining. Surface staining antibody cocktail was added. Cells were fixed with BD CytoFix/CytoPerm for 20 minutes on ice. Cells were permeabilized overnight at 4°C with the FoxP3 Permeabilization kit. After permeabilization, intracellular staining was done in 1X permeabilization buffer (Invitrogen). Cells were washed with permeabilization buffer. Cells were run through the BD LSRII for quantification of cytokine staining. Gating and quantification of populations and mean fluorescent intensity was performed with FlowJov10.8.1.

### Luciferase Assay

Luciferase activity in the cell culture supernatant was quantified using the Nano-Glo® Luciferase Assay System (Promega). 50μL of cell culture supernatant was combined with 50μL of diluted substrate solution and mixed well with a multi-channel pipette. Luciferase activity was measured using the Agilent Bio Tek Synergy neo2 multi-mode plate reader and the Agilent Bio Tek Gen5 software.

### Cell death screen for p35NL release

To determine whether p35NL would be released in response to cell death (Figure 2), 7 x 10^4^ p35NL or NLO retrovirally transduced L cells were exposed to chemical inducers of cell death: actinomycin D, apoptosis activator 2 (Tocris), cycloheximide, oxaliplatin (Tocris), and ionomycin (Tocris). 24 hours after treatment, plates were spun down at 2000 rpm for 7 min at 4°C. Supernatants were drawn off plates and transferred to an empty 96-well round-bottom plate. Supernatants were spun down again to ensure a cell-free supernatant, and 50μL of supernatants were then used for luciferase assay.

### Cell viability measurements

After the removal of supernatant, L cells were trypsinized with 0.25% trypsin and resuspended in complete RPMI 1640 media. Cell viability was determined with 1:1 cell suspension to ViaStain™ AOPI staining (Revvity) and quantified using the Cellaca MX™ cell counter and Matrix software (Revvity).

### Lactate dehydrogenase (LDH) assay for cell viability

LDH release was measured in cells treated with chemical inducers of cell death. LDH in the supernatant from cell death screen (described above) was quantified using the CyQuant LDH assay from ThermoFisher.

### High-throughput screening pharmacological screen

A screen of 114 drugs was sourced from Selleck Chem (Ionomycin was added to the screen from previous results in Figure 2). p35NL L cells were plated using Opentrons 2 pipetting robot and exposed to dilutions (100-0.1µM) of 114 drugs for 24 hours. Supernatants were collected after 24 hours of drug exposure for luciferase assay detection of p35NL release.

### Characterization of ionomycin and CB-5083 as inducers of p35 release

2 x 10^4^ p35NL and NLO cells were plated in a 96-well flat-bottom plate (Corning). Cells were exposed to titrations of ionomycin (Tocris), CB-5083 (Tocris), or vehicle for 24 hours. Supernatants were collected from plates as described above and used for luciferase assay to quantify p35NL and NLO release. Cell viability was determined as described above. To determine the kinetics of p35NL and NLO release, supernatants and cell viability were also collected at 4, 8, 12, and 24 time points. For actinomycin D and cycloheximide experiments, both cell lines were pre-treated with the indicated concentrations of ActD or CHX for 30 min and then stimulated with ionomycin and CB-5083 for 24 hours.

### 2D-Gel Electrophoresis

P35NL L cells were serum starved from 10% to 1.5-0.78% FCS gradually over the course of two weeks. p35NL cells were then stimulated with DMSO (CTRL), 3.12μM ionomycin, or 1.56μM CB-5083 for 24 hours. After drug treatment, supernatants were collected and processed for 2D gel electrophoresis by Applied Biomics.

### Live-cell fluorescent imaging

Reporter cell lines were generated through retroviral transduction of L cells with a combination of fluorescent retroviruses for subcellular compartments. For imaging experiments, 5 x 10^3^ cells were plated on µ-Slide 18 Well chamber slides with ibiTreat coating (Ibidi). Cell lines containing ER-HaloTag® were incubated with the Janelia Fluor® JFX650 HaloTag® ligand (Promega) for 1 hour at 37°C and replaced with fresh media before imaging. Live-cell fluorescent images were captured using the Keyence BZ-X810 fluorescence microscope. All images were captured at 100X. For resting localization of p35-Scarlet, BiP-BFP, Golgi-eGFP, and ER-Halotag fluorophores, pixel intensity and localization of fluorophores was visualized using the RBG profile plot function on ImageJ with a manually drawn region of interest (ROI) line. For experiments involving the trafficking of p35-Scarlet, reporter L cells were exposed to DMSO (vehicle), 3.12µM ionomycin, or 1.56µM CB-5083 for 8-24 hours before imaging on the Keyence. Colocalization of p35-Scarlet, BiP-BFP, and Golgi-eGFP during drug treatment was quantified using the BIOP (Bioimaging and Optics Platform) JaCOP plugin in FIJI. Average Pearson’s coefficient in each experimental condition was calculated using the JaCOP plugin from 20 representative images. Each representative image contained 5-10 cells and Pearson’s coefficient was calculated across each representative image without drawing ROIs. Fluorescence threshold was set using Default settings on the JaCOP plugin.

### Electron microscopy

L cells were treated with DMSO vehicle, 3.12µM ionomycin, 1.56µM CB-5083 for 24 hours in a 6-well plate (Corning). After drug treatment, L cells were fixed on the 6-well plate with 2% paraformaldehyde and 2.5% glutaraldehyde in 0.1M sodium cacodylate buffer containing 2mM calcium chloride. Fixed cells were rinsed with and post fixation 1% osmium tetroxide and 1.5% potassium ferrocyanide in 0.1M sodium cacodylate buffer. Samples were then rinsed with 0.1M sodium acetate buffer and stained *en bloc* overnight with 1% uranyl acetate in 0.1M sodium acetate. After staining, samples were rinsed in 0.1M sodium acetate buffer and once with water. Samples were dehydrated through an increasing ethanol series. Samples were taken through a dilution series of ethanol to epoxy resin (2:1, 1:1, 1:2) using LX112. Samples were incubated for 1 hour in pure resin x 2. Resin was removed from wells, and a thin layer of fresh resin was added before inversion into four Beem capsules. Samples polymerized overnight. Beem capsules were filled with resin to the top and returned to the oven for additional 48 hours of polymerization. Beem capsules were removed from wells. Samples were trimmed from blockfaces and cut into 90nm sections onto copper grids. Samples were imaged on the JEOL 1400 Flash at 80kV on an AMT camera. Sample preparation and imaging was performed by University of Maryland School of Medicine’s and School of Dentistry’s Electron Microscopy Core Imaging Facility (EMCIF) in Baltimore, Maryland.

### UPR reporter and inhibition experiments

To characterize the activation of the UPR response in response to ionomycin and CB-5083, UPR-GFP L cells were generated as described above. 2 x 10^4^ UPR-GFP L cells were exposed to DMSO (vehicle), 3.12µM ionomycin, or 1.56µM CB-5083 for 8, 12, and 24 hours. For each time point, cells were harvested with trypsinization and stained for viability using 7-Aminoactinomycin D. The percentage of UPR+ cells was quantified using flow cytometry on the BD LSR II. UPR+ cells were gated as GFP+7-AAD-cells using FlowJov10.8.1.

The following inhibitors were used to target specific arms of the UPR response: AMG PERK 44 for PERK, MKC8866 for IRE1α, and Ceapin-A7 for ATF6 (Med Chem Express). 2 x 10^4^ p35NL or NLO cells were pre-treated with inhibitors (25-0.39µM) for each of the sensors of the UPR response for 1 hour before the addition of DMSO (vehicle), 3.12µM ionomycin, or 1.56µM CB-5083 for 24 hours. Supernatants were collected as described above and viability was accessed as described above.

### Generation of primary muscle fibroblasts

Muscle tissue was collected from PN3 IL-12p40KO mice. Muscle tissue was dissociated using trypsin and collagenase D digestion and processed into single cell suspensions with mesh. Single-cell suspensions were transferred to a 96-well plate with complete RPMI. After 3 days of differentiation, debris was removed from suspensions and fibroblast morphology was confirmed under the microscope. p40KO muscle fibroblasts were transduced with p35NL or NLO retrovirus as described above. Transduced fibroblasts were treated with titrations of ionomycin and CB-5083 for 24 hours. Supernatants were collected and used for luciferase assays as described above. Viability was determined described above.

### Statistics

Statistics were analyzed using GraphPad Prism 9. Significant differences in relative luminescence units and viability were determined using a one-way ANOVA with Bonferoni’s corrections for multiple comparisons.

## Supporting information

Supplementary Material (Hill et al)

## Supplementary Materials

Supplementary Figure 1. Release of LDH in response to chemical induction of cell death.

Supplementary Figure 2. High-throughput screening of pharmacological triggers for p35 release.

Supplementary Figure 3. Kinetics of p35 release in response to ionomycin stimulation and VCP inhibition.

Supplementary figure 4. Response of p35NL and NLO cells to temporary exposure to ionomycin and CB-5083.

Supplemental Figure 5. 2D-DIGE analysis characterizes the secretome of CB-5083- and ionomycin-treated p35NL L cells.

Supplementary figure 6. Effect of UPR sensor inhibition on p35NL cell viability and NLO release.

## Acknowledgements

We thank the members of the Nevil Singh Lab at UMSOM for vibrant discussions and critique, and the Flow Cytometry Shared Service of the University of Maryland Marlene and Stewart Greenebaum Comprehensive Cancer Center for flow sorting. These studies were supported by National Institutes of Health grant R01AI168192 to NJS.

## Author Contributions

Conceptualization: EMH, NJS

Methodology: EMH, JB, GM, NJS

Investigation: EMH, JB, GM, BC, KR, NJS

Visualization: EMH, BC, NJS

Funding acquisition: NJS Writing: EMH, NJS

Writing – edits and revisions: EMH, JB, GM, BC, KR, NJS

## Data and materials availability

All data are available in the main text or the supplementary materials. Plasmids developed for the study will be available from Addgene upon publication of the manuscript. All code used will be available on http://github.com/NevilLab/

## References

1. K. Abdi, N. J. Singh, Making many from few: IL-12p40 as a model for the combinatorial assembly of heterodimeric cytokines. Cytokine 76, 53–57 (2015).

2. T. S. Kim et al., Epithelial-derived interleukin-23 promotes oral mucosal immunopathology. Immunity 57, 859–875.e811 (2024).

3. P. Matzinger, T. Kamala, Tissue-based class control: the other side of tolerance. Nat Rev Immunol 11, 221–230 (2011).

4. A. N. Gerber, K. Abdi, N. J. Singh, The subunits of IL-12, originating from two distinct cells, can functionally synergize to protect against pathogen dissemination in vivo. Cell Rep 37, 109816 (2021).

5. M. Kobayashi et al., Identification and purification of natural killer cell stimulatory factor (NKSF), a cytokine with multiple biologic effects on human lymphocytes. J Exp Med 170, 827–845 (1989).

6. A. S. Stern et al., Purification to homogeneity and partial characterization of cytotoxic lymphocyte maturation factor from human B-lymphoblastoid cells. Proc Natl Acad Sci U S A 87, 6808–6812 (1990).

7. K. Abdi, IL-12: the role of p40 versus p75. Scandinavian journal of immunology 56, 1–11 (2002).

8. A. D’Andrea et al., Production of natural killer cell stimulatory factor (interleukin 12) by peripheral blood mononuclear cells. J Exp Med 176, 1387–1398 (1992).

9. S. E. Macatonia et al., Dendritic cells produce IL-12 and direct the development of Th1 cells from naive CD4+ T cells. J Immunol 154, 5071–5079 (1995).

10. S. Reitberger, P. Haimerl, I. Aschenbrenner, J. Esser-von Bieren, M. J. Feige, Assembly-induced folding regulates interleukin 12 biogenesis and secretion. The Journal of biological chemistry 292, 8073–8081 (2017).

11. L. W. Collison et al., IL-35-mediated induction of a potent regulatory T cell population. Nature Immunology 11, 1093–1101 (2010).

12. K. O. Dixon, S. W. van der Kooij, D. A. Vignali, C. van Kooten, Human tolerogenic dendritic cells produce IL-35 in the absence of other IL-12 family members. Eur J Immunol 45, 1736–1747 (2015).

13. C. Ye, H. Yano, C. J. Workman, D. A. A. Vignali, Interleukin-35: Structure, Function and Its Impact on Immune-Related Diseases. J Interferon Cytokine Res 41, 391–406 (2021).

14. Y. G. Mideksa et al., A comprehensive set of ER protein disulfide isomerase family members supports the biogenesis of proinflammatory interleukin 12 family cytokines. J Biol Chem 298, 102677 (2022).

15. K. Abdi et al., Free IL-12p40 monomer is a polyfunctional adaptor for generating novel IL-12-like heterodimers extracellularly. J Immunol 192, 6028–6036 (2014).

16. S. Sen Santara et al., The NK cell receptor NKp46 recognizes ecto-calreticulin on ER-stressed cells. Nature 616, 348–356 (2023).

17. M. Obeid et al., Calreticulin exposure dictates the immunogenicity of cancer cell death. Nat Med 13, 54–61 (2007).

18. P. Scaffidi, T. Misteli, M. E. Bianchi, Release of chromatin protein HMGB1 by necrotic cells triggers inflammation. Nature 418, 191–195 (2002).

19. R. Jalah et al., The p40 subunit of interleukin (IL)-12 promotes stabilization and export of the p35 subunit: implications for improved IL-12 cytokine production. The Journal of biological chemistry 288, 6763–6776 (2013).

20. J. Kleeff, M. Kornmann, H. Sawhney, M. Korc, Actinomycin D induces apoptosis and inhibits growth of pancreatic cancer cells. Int J Cancer 86, 399–407 (2000).

21. D. J. Anderson et al., Targeting the AAA ATPase p97 as an Approach to Treat Cancer through Disruption of Protein Homeostasis. Cancer Cell 28, 653–665 (2015).

22. K. Kitakaze et al., Cell-based HTS identifies a chemical chaperone for preventing ER protein aggregation and proteotoxicity. eLife 8, e43302 (2019).

23. I. Pontisso, R. Ornelas-Guevara, E. Chevet, L. Combettes, G. Dupont, Gradual ER calcium depletion induces a progressive and reversible UPR signaling. PNAS Nexus 3, pgae229 (2024).

24. C. Li et al., p97/VCP is highly expressed in the stem-like cells of breast cancer and controls cancer stemness partly through the unfolded protein response. Cell Death & Disease 12, 286 (2021).

25. S. E. Logue et al., Inhibition of IRE1 RNase activity modulates the tumor cell secretome and enhances response to chemotherapy. Nature Communications 9, 3267 (2018).

26. M. McLaughlin et al., Inhibition of secretion of interleukin (IL)-12/IL-23 family cytokines by 4-trifluoromethyl-celecoxib is coupled to degradation via the endoplasmic reticulum stress protein HERP. J Biol Chem 285, 6960–6969 (2010).

27. T. L. Murphy, M. G. Cleveland, P. Kulesza, J. Magram, K. M. Murphy, Regulation of interleukin 12 p40 expression through an NF-kappa B half-site. Mol Cell Biol 15, 5258–5267 (1995).

28. C. Reis e Sousa et al., In vivo microbial stimulation induces rapid CD40 ligand-independent production of interleukin 12 by dendritic cells and their redistribution to T cell areas. J Exp Med 186, 1819–1829 (1997).

29. K. Abdi, N. J. Singh, Antigen-activated T cells induce IL-12p75 production from dendritic cells in an IFN-γ-independent manner. Scand J Immunol 72, 511–521 (2010).

30. C. Liu, T. E. Hermann, Characterization of ionomycin as a calcium ionophore. Journal of Biological Chemistry 253, 5892–5894 (1978).

31. S. Yoshida, S. Plant, Mechanism of release of Ca2+ from intracellular stores in response to ionomycin in oocytes of the frog Xenopus laevis. J Physiol 458, 307–318 (1992).

32. C. Booth, G. L. Koch, Perturbation of cellular calcium induces secretion of luminal ER proteins. Cell 59, 729–737 (1989).

33. M. Asano et al., Endoplasmic reticulum resident, immunoglobulin joining chain, can be secreted by perturbation of the calcium concentration in the endoplasmic reticulum. DNA Cell Biol 23, 403–411 (2004).

34. C. K. Suzuki, J. S. Bonifacino, A. Y. Lin, M. M. Davis, R. D. Klausner, Regulating the retention of T-cell receptor alpha chain variants within the endoplasmic reticulum: Ca(2+)-dependent association with BiP. J Cell Biol 114, 189–205 (1991).

35. J. Shen, X. Chen, L. Hendershot, R. Prywes, ER stress regulation of ATF6 localization by dissociation of BiP/GRP78 binding and unmasking of Golgi localization signals. Dev Cell 3, 99–111 (2002).

36. A. Bertolotti, Y. Zhang, L. M. Hendershot, H. P. Harding, D. Ron, Dynamic interaction of BiP and ER stress transducers in the unfolded-protein response. Nat Cell Biol 2, 326–332 (2000).

37. T. Yoneda et al., Activation of caspase-12, an endoplastic reticulum (ER) resident caspase, through tumor necrosis factor receptor-associated factor 2-dependent mechanism in response to the ER stress. J Biol Chem 276, 13935–13940 (2001).

38. A. B. Tam et al., The UPR Activator ATF6 Responds to Proteotoxic and Lipotoxic Stress by Distinct Mechanisms. Developmental Cell 46, 327–343.e327 (2018).

39. J. Sargeant et al., ALG-2 and peflin regulate COPII targeting and secretion in response to calcium signaling. J Biol Chem 297, 101393 (2021).

40. L. Stach, P. S. Freemont, The AAA+ ATPase p97, a cellular multitool. Biochem J 474, 2953–2976 (2017).

41. S. Kilgas, K. Ramadan, Inhibitors of the ATPase p97/VCP: From basic research to clinical applications. Cell Chemical Biology 30, 3–21 (2023).

42. W. K. Xu, Y. Gou, M. M. Lozano, J. P. Dudley, Unconventional p97/VCP-Mediated Endoplasmic Reticulum-to-Endosome Trafficking of a Retroviral Protein. J Virol 95, e0053121 (2021).

43. E. Kim, D. M. Lee, M. J. Seo, H. J. Lee, K. S. Choi, Intracellular Ca(2 +) Imbalance Critically Contributes to Paraptosis. Front Cell Dev Biol 8, 607844 (2020).

44. B. Monel et al., Zika virus induces massive cytoplasmic vacuolization and paraptosis-like death in infected cells. Embo j 36, 1653–1668 (2017).

45. S. Gil-Parrado et al., Ionomycin-activated calpain triggers apoptosis. A probable role for Bcl-2 family members. J Biol Chem 277, 27217–27226 (2002).

46. Z. Shao et al., The protein segregase VCP/p97 promotes host antifungal defense via regulation of SYK activation. PLoS Pathog 20, e1012674 (2024).

47. S. Ghosh et al., Host AAA-ATPase VCP/p97 lyses ubiquitinated intracellular bacteria as an innate antimicrobial defence. Nature Microbiology 10, 1099–1114 (2025).

48. S. Dramsi, P. Cossart, Listeriolysin O-mediated calcium influx potentiates entry of Listeria monocytogenes into the human Hep-2 epithelial cell line. Infect Immun 71, 3614–3618 (2003).

49. S. Roy, S. Roy, S. Halder, K. Jana, A. Ukil, Leishmania exploits host cAMP/EPAC/calcineurin signaling to induce an IL-33-mediated anti-inflammatory environment for the establishment of infection. J Biol Chem 300, 107366 (2024).

50. F. Wang et al., Targeting VCP potentiates immune checkpoint therapy for colorectal cancer. Cell Reports 42, (2023).

