## Supplementary Material (Hill et al) for "The active secretion of a subunit of IL-12 by tissue cells is regulated by Valosin-Containing Protein and intracellular calcium redistribution"

List of Supplemental Figures:

Supplementary Figure 1. Release of LDH in response to chemical induction of cell death.

Supplementary Figure 2. High-throughput screening of pharmacological triggers for p35 release.

Supplementary Figure 3. Kinetics of p35 release in response to ionomycin stimulation and VCP inhibition.

Supplementary figure 4. Response of p35NL and NLO cells to temporary exposure to ionomycin and CB-5083.

Supplemental Figure 5. 2D-DIGE analysis characterizes the secretome of CB-5083- and ionomycin-treated p35NL L cells.

Supplementary figure 6. Effect of UPR sensor inhibition on p35NL cell viability and NLO release.

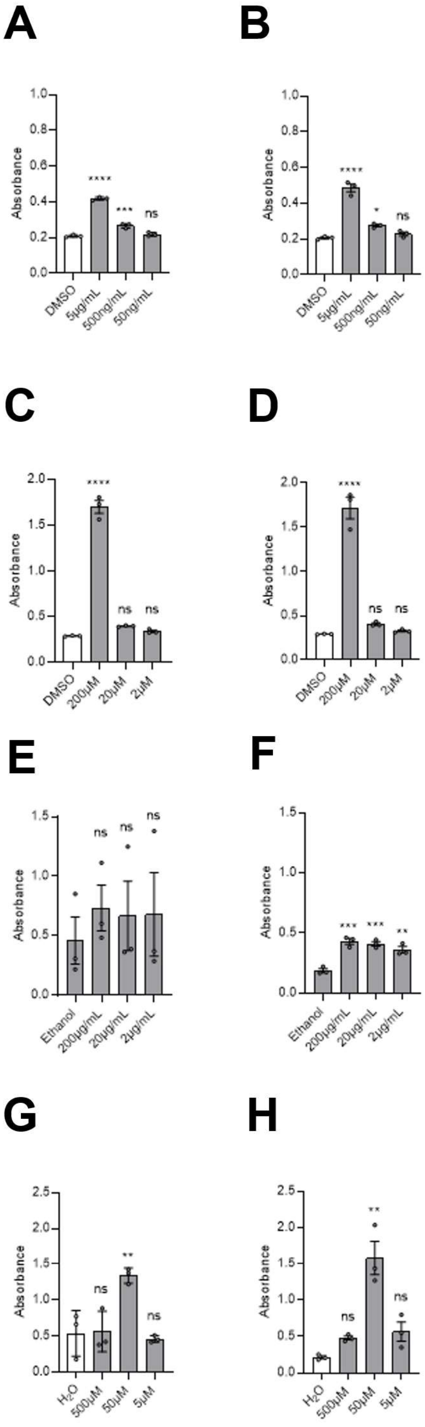
A
B
C
D
E
F
G
H
Absorbance
DMSO
5µg/mL
500ng/mL
50ng/mL
200µM
20µM
2µM
Ethanol
200µg/mL
20µg/mL
2µg/mL
H2O
500µM
50µM
5µM
ns

**Supplementary Figure 1. Release of LDH in response to chemical induction of cell death.**

Murine L cells were retrovirally transduced with p35NL or NLO constructs. Cells were exposed to a variety of chemical inducers of cell death. Release of LDH into the cell culture supernatant was quantified using the CyQuant LDH Cytotoxicity Assay (Invitrogen).

LDH assay of supernatants from p35NL cells exposed to (A) actinomycin D, (C) apoptosis activator 2, (E) cycloheximide, and (G) oxaliplatin.

LDH assay of supernatants from NLO L cells exposed to (B) actinomycin D, (D) apoptosis activator 2, (F) cycloheximide, and (H) oxaliplatin.

Data is represented as the mean corrected absorbance corresponding to LDH quantification (n = 3). Significant changes in LDH in the cell culture media were determined through one-way ANOVA of the mean in comparison to vehicle controls (DMSO).

*, $p < 0.05$; ** $p < 0.01$; *** $p < 0.001$; **** $p < 0.0001$; ns = not significant

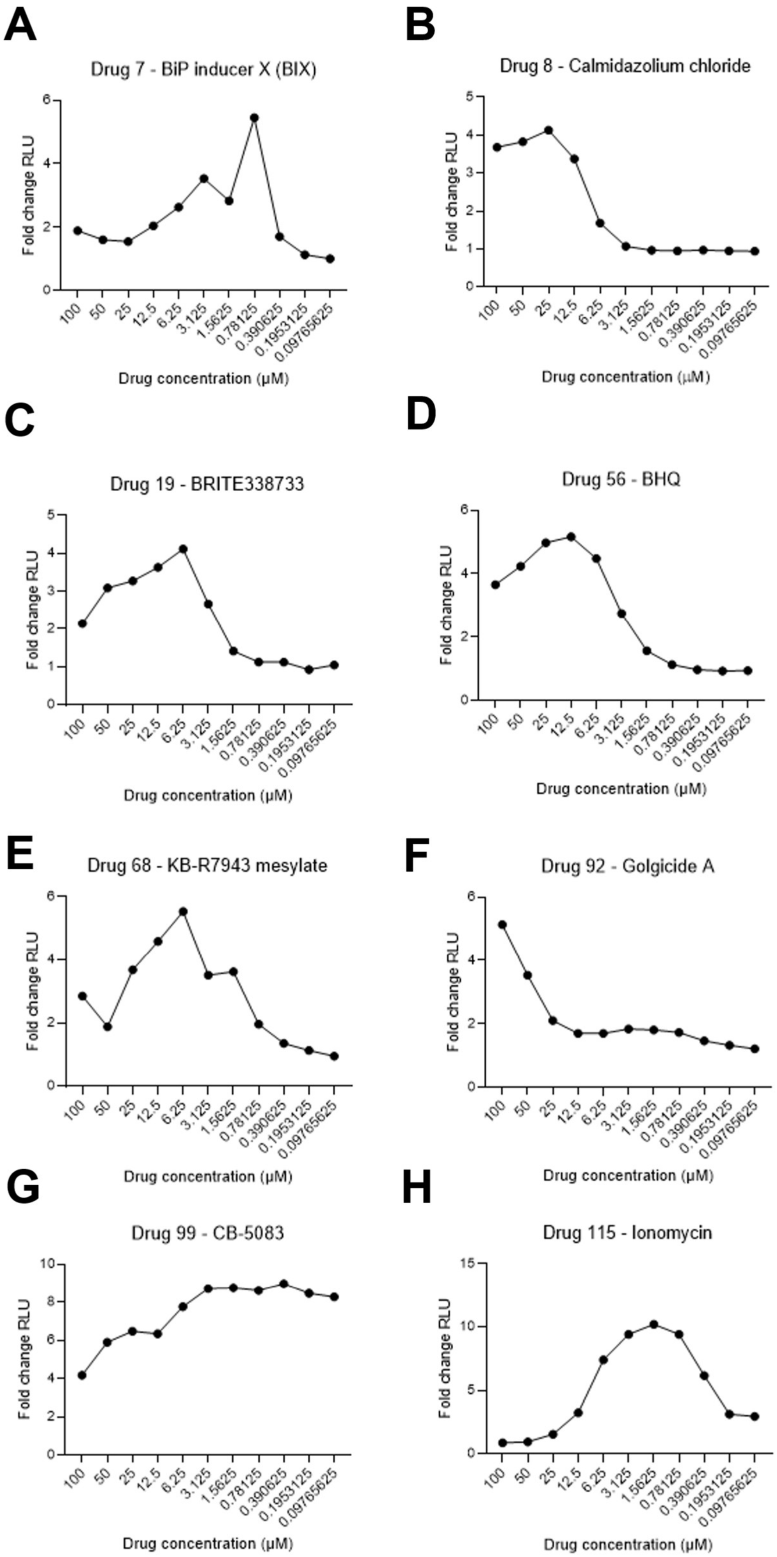
A
Drug 7 - BiP inducer X (BIX)
B
Drug 8 - Calmidazolium chloride
C
Drug 19 - BRITE338733
D
Drug 56 - BHQ
E
Drug 68 - KB-R7943 mesylate
F
Drug 92 - Golgicide A
G
Drug 99 - CB-5083
H
Drug 115 - Ionomycin
Fold change RLU
Drug concentration (μM)
100
50
25
12.5
6.25
3.125
1.5625
0.78125
0.390625
0.1953125
0.09765625

**Supplementary Figure 2. High-throughput screening of pharmacological triggers for p35 release.**

p35-NL L cells were plated using the Opentrons pipetting robot. 115 drugs from the Cherry Select panel were diluted using the Opentrons pipetting robot. p35NL cells were treated with dilutions of drug for 24 hours. Supernatants were harvested and used for luciferase assay to detect p35 release. The following drugs were shown to induce the release of p35 using the cutoff of a fold change in RLU in comparison to media only control greater than or equal to 4.

(A) Drug 7 BiP inducer X

(B) Drug 8 Calmidazolium chloride

(C) Drug 19 BRITE338733

(D) Drug 56 BHQ

(E) Drug 68 KB-R7943 mesylate

(F) Drug 92 Golgicide A

(G) Drug 99 CB-5083

(H) Drug 115: Ionomycin

Data is represented as the mean of two independent plates normalized to media-only controls. (n = 2).

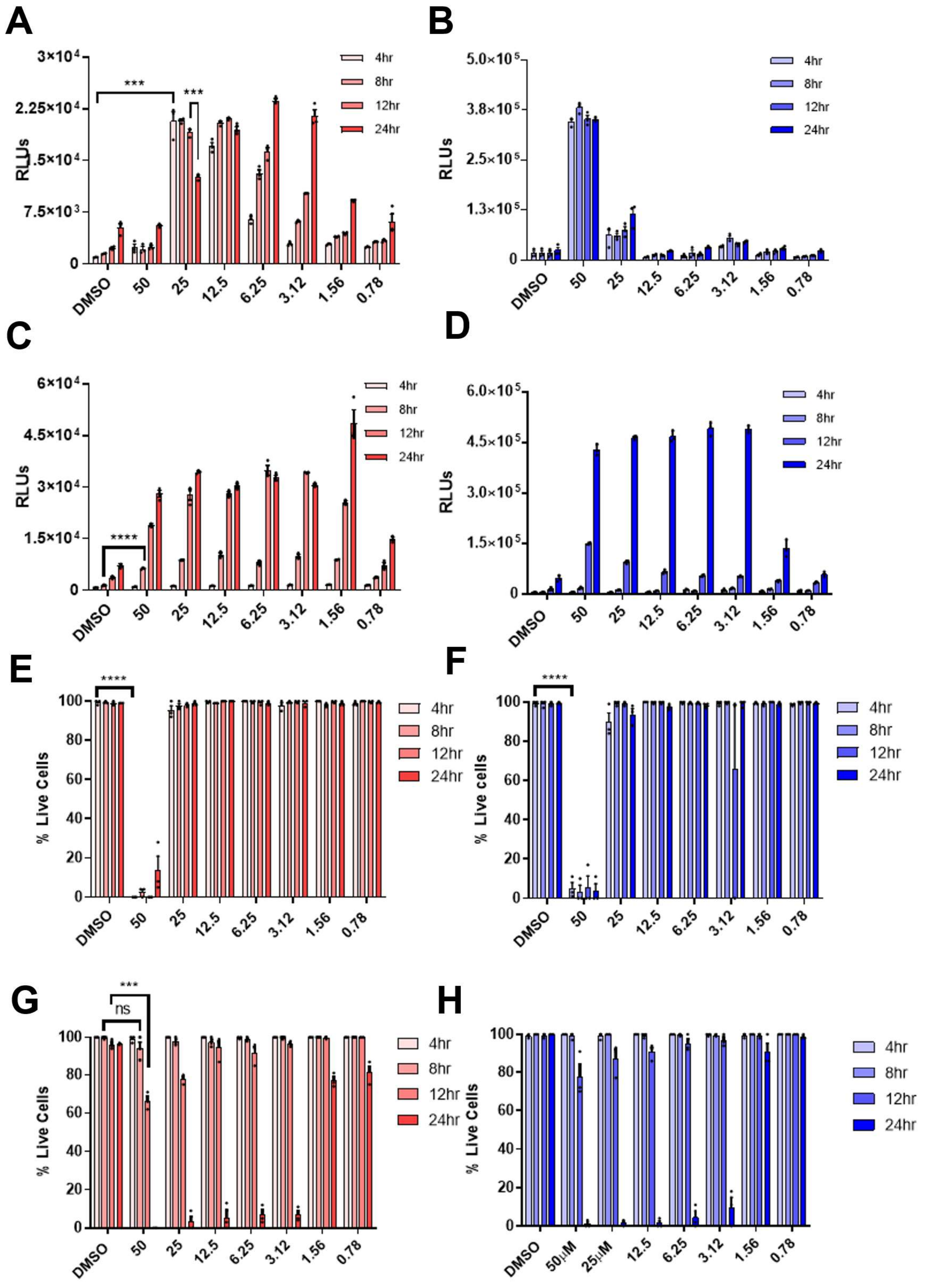

A
B
C
D
E
F
G
H
RLUs
% Live Cells
% Live cells
4hr
8hr
12hr
24hr
DMSO
50
25
12.5
6.25
3.12
1.56
0.78
50μM
25μM
***
****
ns

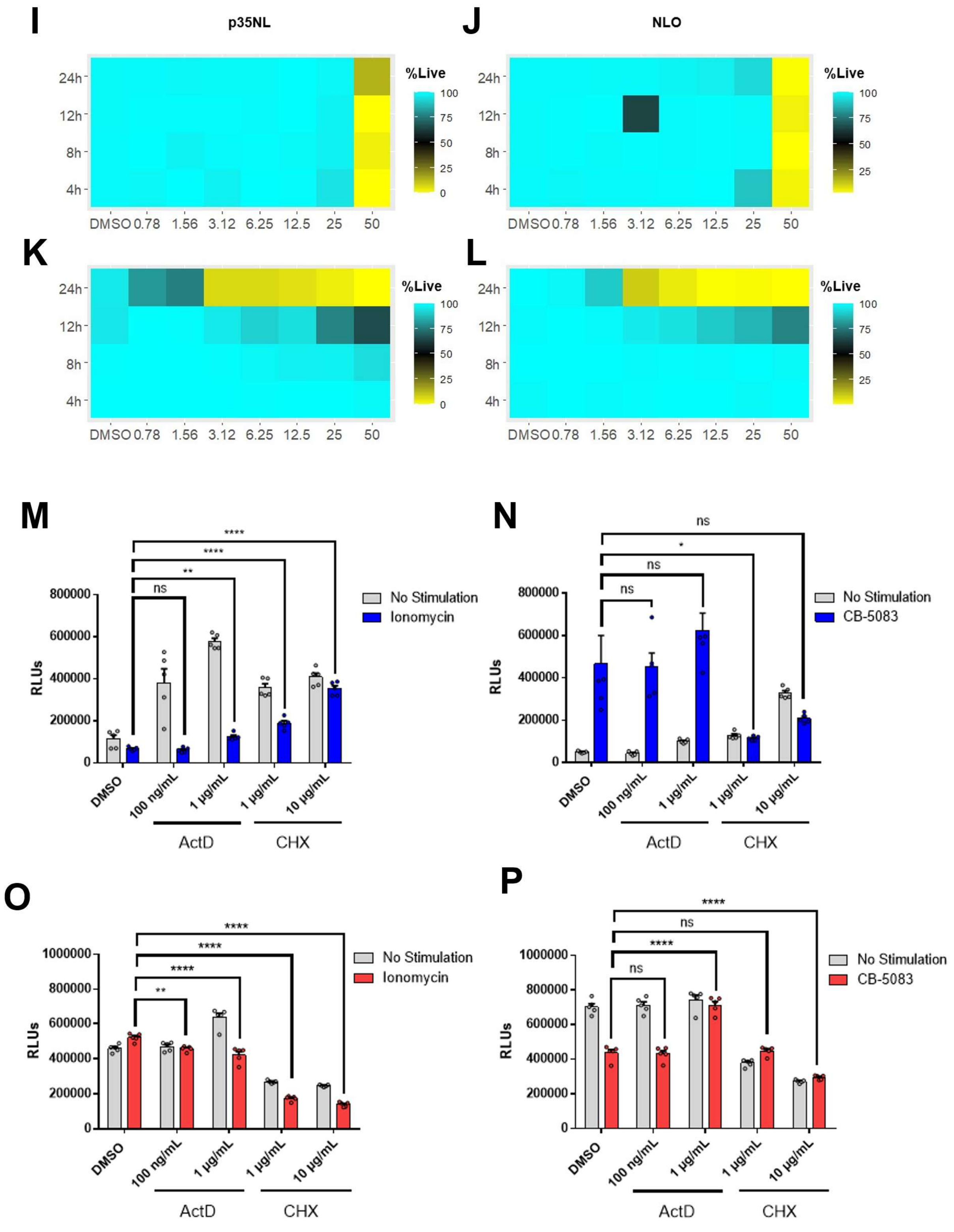
I
p35NL
J
NLO
%Live
100
75
50
25
0
24h
12h
8h
4h
DMSO 0.78 1.56 3.12 6.25 12.5 25 50
K
L
M
N
RLUs
No Stimulation
Ionomycin
CB-5083
DMSO
100 ng/mL
1 μg/mL
1 μg/mL
10 μg/mL
ActD
CHX
ns
*
**
****
O
P
1000000
800000
600000
400000
200000
0

**Supplementary Figure 3. Kinetics of p35 release in response to ionomycin stimulation and VCP inhibition.**

(A) Luciferase assay showing the kinetics of p35NL release in response to ionomycin stimulation at 4-, 8-, 12-, and 24-hour time points. (B) Luciferase assay showing the kinetics of NLO release in response to ionomycin stimulation at 4-, 8-, 12-, and 24-hour time points. Viability of p35NL cells exposed to ionomycin for 4-, 8-, 12-, and 24-hour timepoints. (C) Luciferase assay showing the kinetics of p35NL release in response to CB-5083 treatment at 4-, 8-, 12-, and 24-hour time points. (D) Luciferase assay showing the kinetics of NLO release in response to CB-5083 treatment at 4-, 8-, 12-, and 24-hour time points. (E) Viability of p35NL cells exposed to ionomycin for 4-, 8-, 12-, and 24-hour timepoints. (F) Viability of NLO cells exposed to ionomycin for 4-, 8-, 12-, and 24-hour timepoints. (G) Viability of p35NL cells exposed to CB-5083 for 4-, 8-, 12-, and 24-hour timepoints. (H) Viability of NLO cells exposed to CB-5083 for 4-, 8-, 12-, and 24-hour timepoints. (n=3)

(I) Heat map of cell viability showing the kinetics of cell death in p35NL cells in response to ionomycin. (J) Heat map of cell viability showing the kinetics of cell death in NLO cells in response to ionomycin. (K) Heat map of cell viability showing the kinetics of cell death in p35NL cells in response to CB-5083. (L) Heat map of cell viability showing the kinetics of cell death in NLO cells in response to CB-5083 (n=3).

(M) NLO cells were pre-treated with DMSO, actinomycin D, or cycloheximide for 30 minutes before the addition of vehicle or 6.25µM ionomycin. Supernatants were collected after 24 hours and used for luciferase assay. (N) NLO cells were pre-treated with DMSO, actinomycin D, or cycloheximide for 30 minutes before the addition of vehicle or 1.56µM CB-5083. Supernatants were collected after 24 hours and used for luciferase assay. (O) Intracellular p35NL measurements from cells pre-treated with DMSO, actinomycin D, or cycloheximide for 30 minutes before the addition of vehicle or 6.25µM ionomycin for 24 hours (P) Intracellular p35NL measurements from cells pre-treated with DMSO, actinomycin D, or cycloheximide for 30 minutes before the addition of vehicle or 1.56µM CB-5083 for 24 hours.

Data is represented as raw luminescence units and the percentage of live cells quantified from propidium iodide.

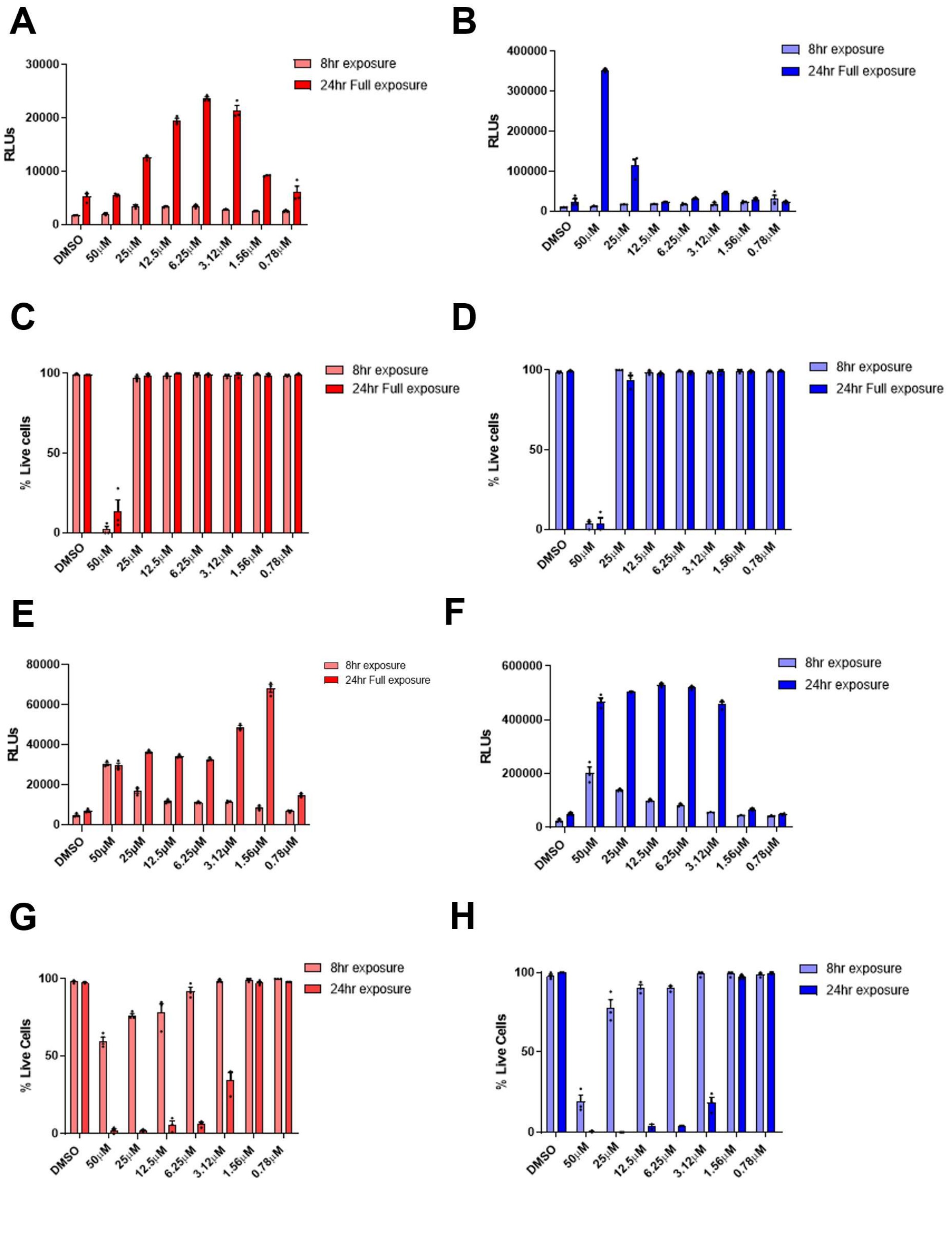
A
8hr exposure
24hr Full exposure
RLUs
30000
20000
10000
0
DMSO
50μM
25μM
12.5μM
6.25μM
3.12μM
1.56μM
0.78μM
B
8hr exposure
24hr Full exposure
RLUs
400000
300000
200000
100000
0
C
8hr exposure
24hr Full exposure
% Live cells
100
50
0
D
8hr exposure
24hr Full exposure
% Live cells
E
8hr exposure
24hr Full exposure
RLUs
80000
60000
40000
20000
0
F
8hr exposure
24hr exposure
RLUs
600000
400000
200000
0
G
8hr exposure
24hr exposure
% Live Cells
H
8hr exposure
24hr exposure
% Live Cells

**Supplementary figure 4. Response of p35NL and NLO cells to temporary exposure to ionomycin and CB-5083.**

(A) Luciferase assay comparing supernatants from p35NL cells exposed to ionomycin for 8 hours with 16 hours to recover in fresh media compared to p35NL cells exposed to ionomycin for a full 24 hours. (B) Luciferase assay comparing supernatants from NLO cells exposed to ionomycin for 8 hours with 16 hours to recover in fresh media compared to p35NL cells exposed to ionomycin for a full 24 hours (C) Survival of p35NL cells exposed to ionomycin for 8 hours with 16 hours to recover in fresh media compared to survival of p35NL cells exposed to ionomycin for a full 24 hours. (D) Survival of NLO cells exposed to ionomycin for 8 hours with 16 hours to recover in fresh media compared to survival of p35NL cells exposed to ionomycin for a full 24 hours. (E) Luciferase assay comparing supernatants from p35NL cells exposed to CB-5083 for 8 hours with 16 hours to recover in fresh media compared to p35NL cells exposed to CB-5083 for a full 24 hours. (F) Luciferase assay comparing supernatants from NLO cells exposed to ionomycin for 8 hours with 16 hours to recover in fresh media compared to p35NL cells exposed to CB-5083 for a full 24 hours (G) Survival of p35NL cells exposed to CB-5083 for 8 hours with 16 hours to recover in fresh media compared to survival of p35NL cells exposed to CB-5083 for a full 24 hours. (H) Survival of NLO cells exposed to CB-5083 for 8 hours with 16 hours to recover in fresh media compared to survival of p35NL cells exposed to CB-5083 for a full 24 hours.

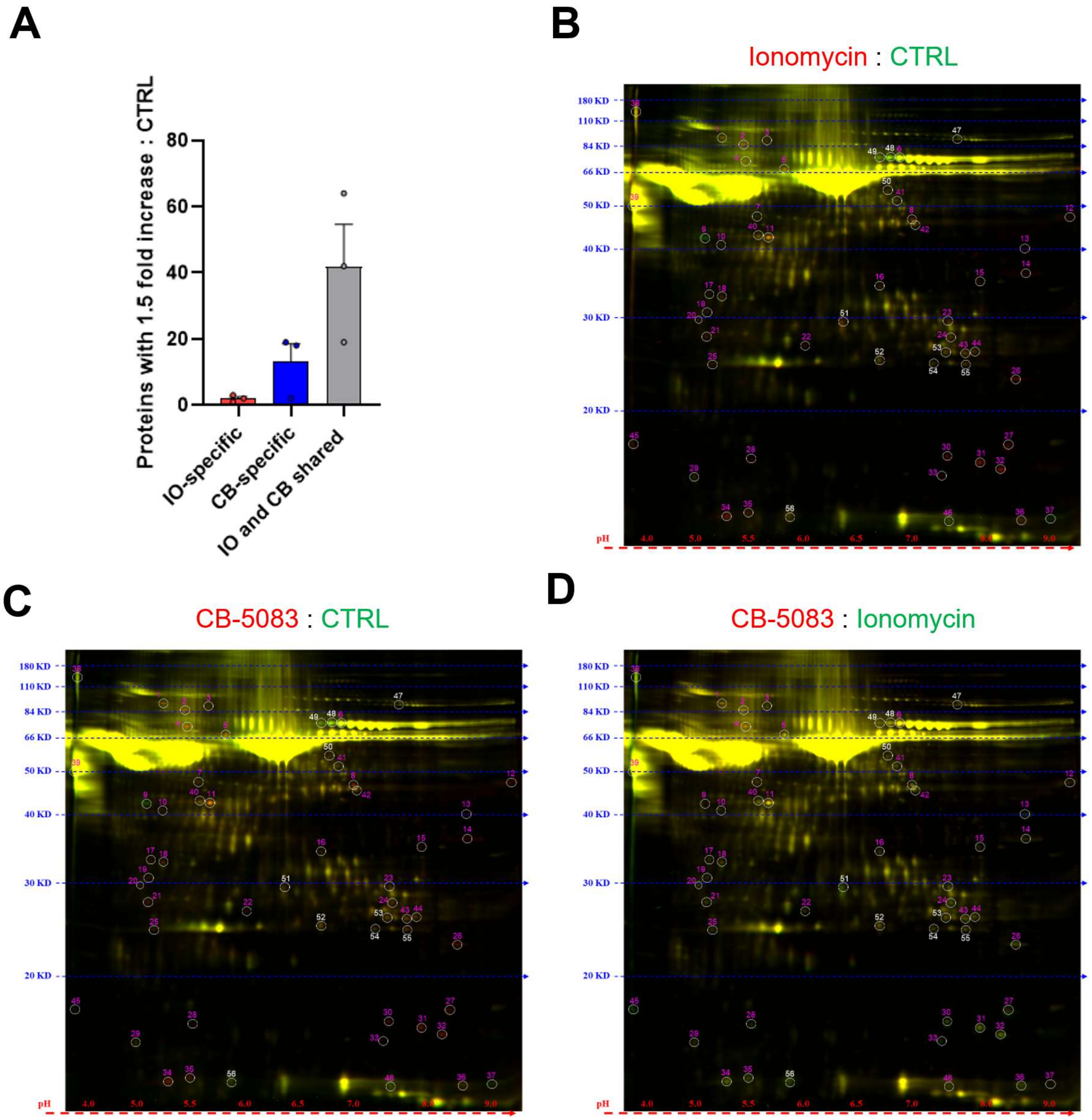

A
B
C
D
Proteins with 1.5 fold increase : CTRL
80
60
40
20
0
IO-specific
CB-specific
IO and CB shared
Ionomycin : CTRL
CB-5083 : CTRL
CB-5083 : Ionomycin
180 KD
110 KD
84 KD
66 KD
50 KD
40 KD
30 KD
20 KD
pH
4.0
5.0
5.5
6.0
6.5
7.0
9.0

Supplemental Figure 5. 2D-DIGE analysis characterizes the “secretome” of CB-5083- and ionomycin-treated p35NL L cells.

p35NL L cells were stimulated with either DMSO (CTRL), 3.12µM ionomycin, or 1.56µM CB-5083 for 24 hours. Secretome was analyzed using 2D-DIGE analysis. (A) Comparison of released proteins from the supernatant of ionomycin (IO) and CB-5083-treated (CB) p35NL cells. A cut-off of 1.5-fold increase in comparison to vehicle controls was used. Results are pooled from three separate gels in which cells were serum starved to 1.5% or 0.78% FCS. (B) Representative 2D gel comparing the proteins from CTRL supernatant (green) to ionomycin-treated supernatants (red). (C) Representative 2D gel comparing the proteins from CTRL supernatants (green) to CB-5083-treated supernatants (red). (D) Representative 2D gel comparing the proteins from ionomycin-treated supernatants (green) to CB-5083-treated supernatants (red).

**Supplementary figure 6. Effect of UPR sensor inhibition on p35NL cell viability and NLO release:**

(A) Luciferase assay from p35NL cells exposed to the PERK inhibitor AMG PERK 44 for 24 hours. (B) Viability of p35NL cells exposed to AMG PERK 44 for 24 hours. (C) Viability of p35NL cells pre-treated with AMG PERK 44 for 1 hour and stimulated with 3.12µM ionomycin for 24 hours. (D) Viability of p35NL cells pre-treated with AMG PERK 44 for 1 hour and stimulated with 1.56µM CB-5083 for 24 hours. (E) Luciferase assay of NLO cells treated exposed to the PERK inhibitor AMG PERK 44 for 24 hours. (F) Luciferase assay of NLO cells pre-treated with AMG PERK 44 for 1 hour and stimulation with 3.12µM ionomycin for 24 hours. (G) Luciferase assay of NLO cells pre-treated with AMG PERK 44 for 1 hour and stimulation with 1.56µM CB-5083 for 24 hours. (H) Viability of NLO cells exposed to AMG PERK 44 for 24 hours. (I) Viability of NLO cells pre-treated with AMG PERK 44 for 1 hour and stimulated with 3.12µM ionomycin for 24 hours. (J) Viability of NLO cells pre-treated with AMG PERK 44 for 1 hour and stimulated with 1.56µM CB-5083 for 24 hours.

(K) Luciferase assay from p35NL cells exposed to the IRE1α inhibitor MKC8866 for 24 hours. (L) Viability of p35NL cells exposed to MKC8866 for 24 hours. (M) Viability of p35NL cells pre-treated with MKC8866 for 1 hour and stimulated with 3.12µM ionomycin for 24 hours. (N) Viability of p35NL cells pre-treated with MKC8866 for 1 hour and stimulated with 1.56µM CB-5083 for 24 hours. (O) Luciferase assay of NLO cells treated exposed to the IRE1α inhibitor MKC8866 for 24 hours. (P) Luciferase assay of NLO cells pre-treated with MKC8866 for 1 hour and stimulation with 3.12µM ionomycin for 24 hours. (Q) Luciferase assay of NLO cells pre-treated with MKC8866 for 1 hour and stimulation with 1.56µM CB-5083 for 24 hours. (R) Viability of NLO cells exposed to MKC8866 for 24 hours. (S) Viability of NLO cells pre-treated with MKC8866 for 1 hour and stimulated with 3.12µM ionomycin for 24 hours. (T) Viability of NLO cells pre-treated with MKC8866 for 1 hour and stimulated with 1.56µM CB-5083 for 24 hours.

(U) Luciferase assay from p35NL cells exposed to the ATF6 inhibitor Ceapin-A7 for 24 hours. (W) Viability of p35NL cells exposed to Ceapin-A7 for 24 hours. (X) Viability of p35NL cells pre-treated with Ceapin-A7 for 1 hour and stimulated with 3.12µM ionomycin for 24 hours. (Y) Viability of p35NL cells pre-treated with Ceapin-A7 for 1 hour and stimulated with 1.56µM CB-5083 for 24 hours. (Z) Luciferase assay of NLO cells treated exposed to the ATF6 inhibitor Ceapin-A7 for 24 hours. (AA) Luciferase assay of NLO cells pre-treated with Ceapin-A7 for 1 hour and stimulated with 3.12µM ionomycin for 24 hours. (BB) Luciferase assay of NLO cells pre-treated with Ceapin-A7 for 1 hour and stimulation with 1.56µM CB-5083 for 24 hours. (CC) Viability of NLO cells exposed to Ceapin-A7 for 24 hours. (DD) Viability of NLO cells pre-treated with Ceapin-A7 for 1 hour and stimulated with 3.12µM ionomycin for 24 hours. (EE) Viability of NLO cells pre-treated with Ceapin-A7 for 1 hour and stimulated with 1.56µM CB-5083 for 24 hours.

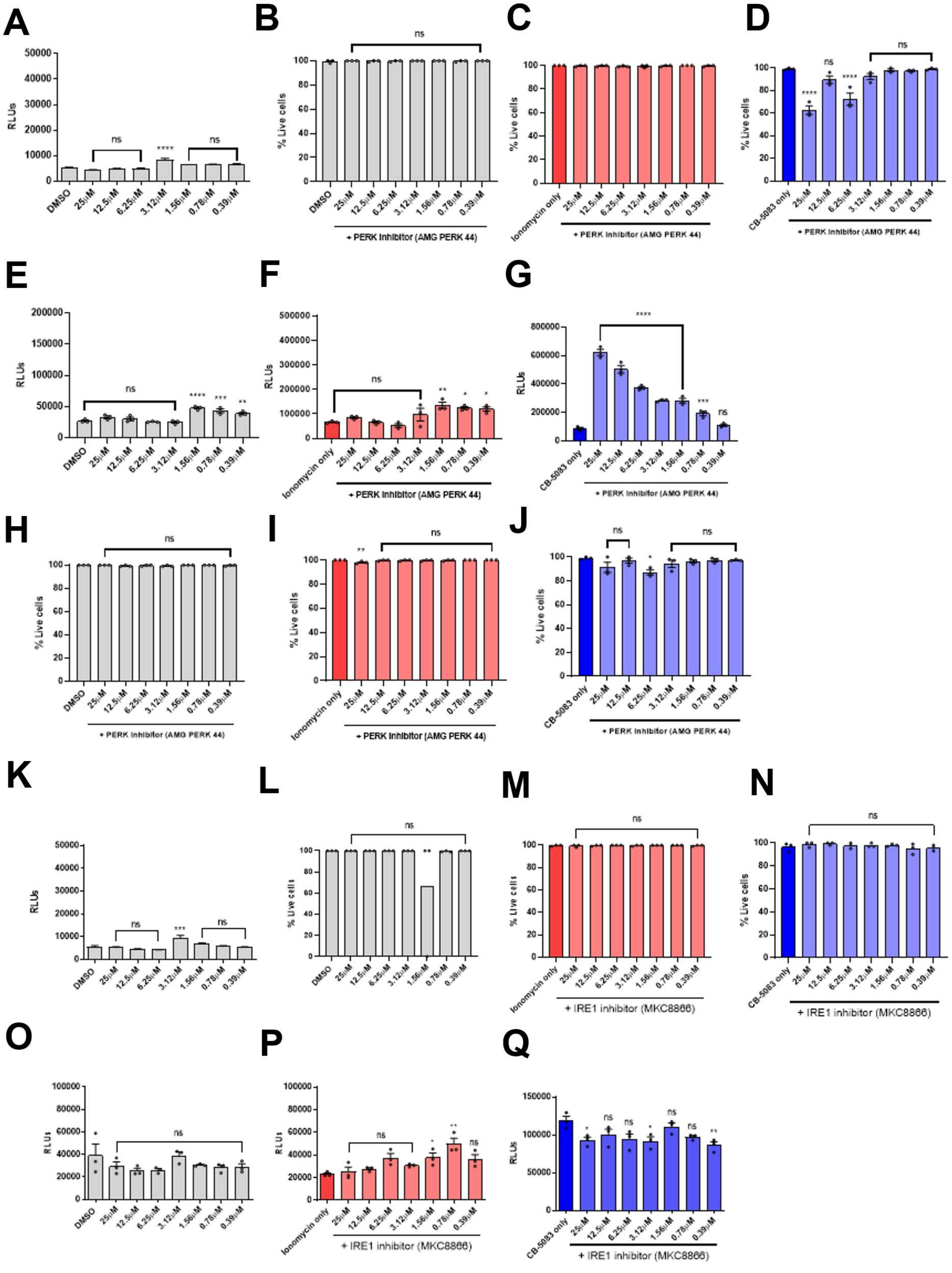

A
RLUs
DMSO
25μM
12.5μM
6.25μM
3.12μM
1.56μM
0.78μM
0.39μM
ns
****
B
% Live cells
+ PERK inhibitor (AMG PERK 44)
C
Ionomycin only
+ PERK inhibitor (AMG PERK 44)
D
CB-5083 only
+ PERK inhibitor (AMG PERK 44)
E
F
G
H
I
J
K
L
M
+ IRE1 inhibitor (MKC8866)
N
O
P
Q

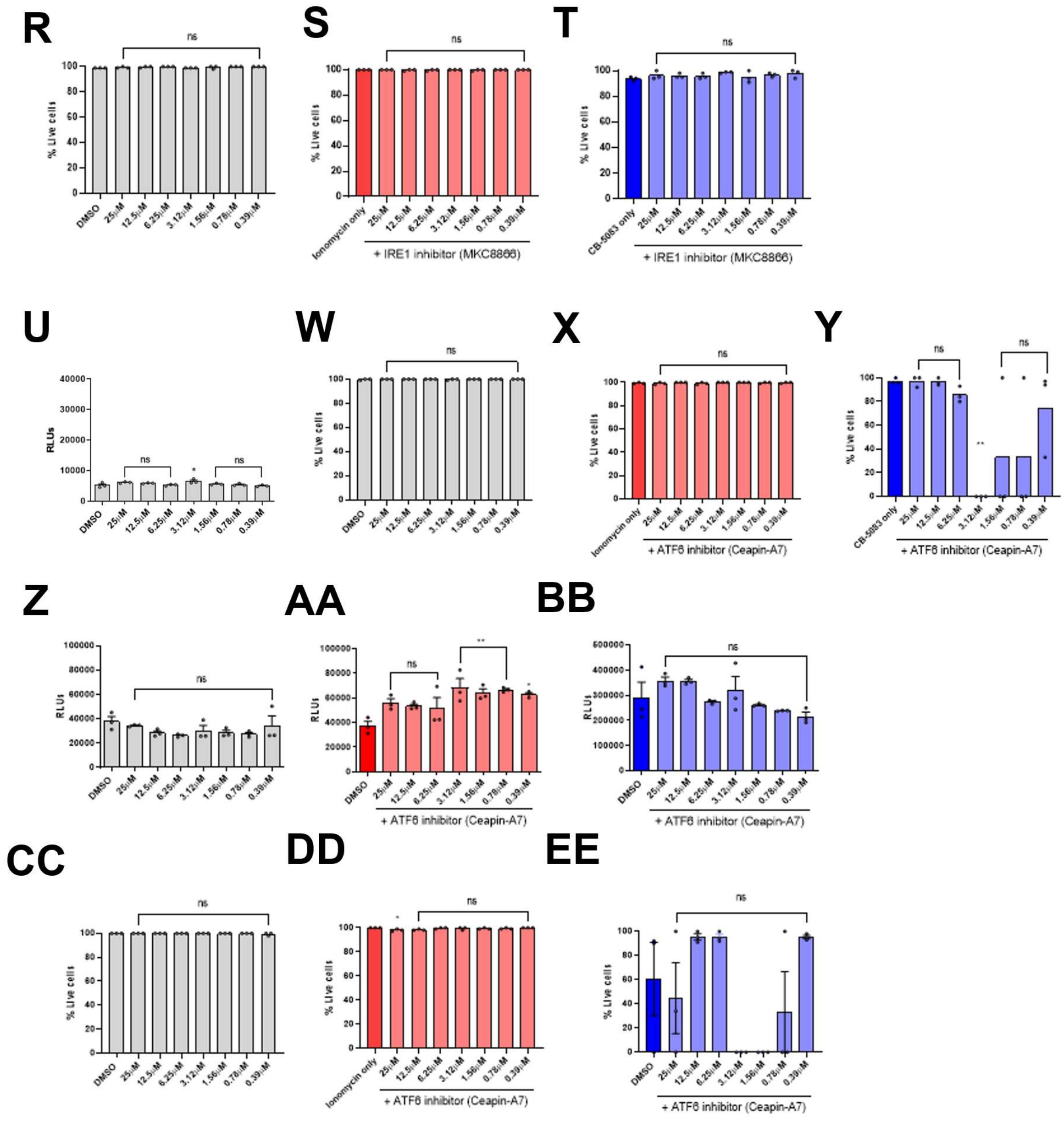
R
S
T
% Live cells
ns
DMSO
25μM
12.5μM
6.25μM
3.12μM
1.56μM
0.78μM
0.39μM
Ionomycin only
CB-5083 only
+ IRE1 inhibitor (MKC8866)
U
W
X
Y
RLUs
+ ATF6 inhibitor (Ceapin-A7)
Z
AA
BB
CC
DD
EE
